# Clinical Implications of SRPK1 Expression in Human Tumours: A Comprehensive Pan-Cancer Analysis Based on Multi-Omics Databases

**DOI:** 10.1101/2025.11.25.690492

**Authors:** Duygu Duzgun, Sebastian Oltean

## Abstract

Serine/arginine-rich protein kinase 1 (SRPK1), which primarily regulates alternative splicing (AS), has been implicated in various malignancies. However, the comprehensive expression landscape and clinical relevance of SRPK1 across diverse tumour types have not been systematically investigated. Chemoresistance remains a formidable obstacle in cancer treatment, accounting for nearly 90% of treatment failures and leading to poor patient survival. Aberrant AS, often driven by splicing factors like SRPK1, is a mechanism cancer cells exploit to overcome chemotherapy-induced cytotoxicity. With that, this study tested the hypothesis of tumour-specific alterations in SRPK1 expression as novel therapeutic targets through a pan-cancer analysis of SRPK1 using multi-omics data, obtained from public databases, including TCGA, CPTAC, GEPIA2, and drug sensitivity platforms. The analysis specifically assessed the pan-cancer expression landscape of SRPK1 at the mRNA and protein levels and its prognostic implications across different cancer types (breast, colon, and prostate cancers), functional analysis of SRPK1-associated splicing network, and its expression in correlation with predicted drug sensitivity. The results revealed the correlations of high SRPK1 expression with worse overall and disease-free survival in a cancer-specific manner and resistance to standard-of-care agents like cisplatin and docetaxel. The integrative multi-omics approach in this study provided a robust foundation for understanding the complex, multifaceted role of SRPK1 in tumour-specific chemoresistance.

## 1. Introduction

As the second leading factor of mortality globally, cancer represents a significant burden on both current and future global health *(Nagai and Kim, 2017)*. In 2022, there were about 20 million new cancer cases and 9.7 million cancer-related death cases, as published in the International Agency for Research on Cancer’s (IARC) GLOBOCAN report *(Bray et al., 2024).* By 2040, the number of cancer-related cases is projected to increase to approximately 28.4 million cases, an increase from 2022 by 42% *(Brenner et al., 2023)*.

In these recent decades, there have been significant advancements in cancer biology research and the development of novel therapeutic approaches; however, the limitations of the treatment have made curing cancer a persevering challenge *(Liu et al., 2024; Samantaray et al., 2024)*. The complex, dynamic, and adaptable nature of cancer must be comprehensively grasped for a more effective management of cancer. In cancer management, resistance to chemotherapy, the first-line treatment, is the biggest drawback of cancer treatment, posing an insurmountable problem *(Cree and Charlton, 2017)*. Chemoresistance, which accounts for nearly 90% of treatment failures, poses a major obstacle in maximising treatment efficiency and is a leading cause of relapse and poor patient survival *(Ramos, Sadeghi, and Tabatabaeian, 2021; Madukwe, 2023)*. However, key mechanisms underlying chemoresistance reflect its profound biological complexity, which requires a multidimensional approach *(Duzgun and Oltean, 2025)*.

Given the intricacies of chemoresistance, **a pan-cancer expression analysis** of any potential genes implicated in chemoresistance is necessary to examine their connection to clinical outcomes and explore the novel molecular mechanisms of chemoresistance. Such approach potentially bolsters the effectiveness of cancer treatment, yielding more robust treatment responses and clinical outcomes.

### 1.1. A Multi-Omics Approach in Cancer Research

High-throughput sequencing technologies have shown tremendous advancements and substantially revolutionised the overall environment and context of cancer research through their resilient and effective methodologies that yield novel possibilities in the study of molecular mechanisms associated with therapy resistance *(Hutter and Zenklusen, 2018; Jiang et al., 2022)*. Central to this endeavour is the adoption of pan-cancer analysis, which is a robust integrative approach with the capacity to comparatively analyse genomic, epigenomic, transcriptomic, and proteomic data across a spectrum of cancer types *(Campbell et al., 2020; Qian et al., 2020)*. There are clinical and multi-omics information from more than 20,000 specimens of 33 tumour types matched with normal tissues available in the Cancer Genome Atlas (TCGA) *(Fonseca-Montaño et al., 2022; Avci et al., 2025)*. Analysis of such a rich dataset helps in establishing connections between gene expression characteristics in pretreatment tumour samples and identifying clinically relevant targets for more robust strategies to effectively deal with cancer resistance.

Recent studies have demonstrated that comparing multi-omics and experimental data can provide a unique opportunity for performing integrative analysis of therapy responders and non-responders *(Vokes et al., 2022; Yang et al., 2022)*. In a multi-omics study on the role of solute carrier family 31 member 1 (SLC31A1), a key cisplatin transporter, in driving resistance to cisplatin, *Qi et al. (2023)* reported lower expression level of SLC31A1 in tumour tissues of renal cell carcinoma, lung adenocarcinoma, and prostate adenocarcinoma than that of their corresponding normal tissues, which was associated with reduced overall survival and progression-free survival in patients *(Qi et al., 2023)*. Additionally, zinc-finger protein 711 (ZNF711) has been reported to enhance SLC31A1 expression through histone methylation. The restored sensitivity to cisplatin in ovarian cancer cell lines through the overexpression of ZNF711 regulates SLC31A1, providing a specific target for cisplatin resistance *(Wu et al., 2021)*. These experimental studies collectively demonstrated the integration of multi-omics data at the pan-cancer level as a robust approach for targeting drug resistance-associated genes and their underlying mechanisms, consequently laying the groundwork for future research on chemoresistance.

Despite numerous studies on this subject, the mechanisms underlying chemoresistance have remained underexplored, especially in cases that involve mutations, gene expression changes, alternative splicing, and post-translational protein modification due to the presence of complex and frequently multifactorial causes *(Aleksakhina, Kashyap, and Imyanitov, 2019)*. With the recent <u>RNA sequencing</u> advancement, a growing body of evidence supporting the notion that drug resistance develops when cancer cells manage to subdue the cytotoxic effects associated with the administered chemotherapeutic agents by **alternative splicing** *(Mehterov et al., 2021)*. Although research on this notion is still in its early stage, elucidating how key regulator-driven splicing networks converge on key pathways to drive chemoresistance yields valuable information on the therapeutic management of cancer. Building on this understanding, the analysis of multi-omics data to identify potential candidates that specifically target abnormal alternative splicing-driven chemoresistance paves the way for future cancer research on drug resistance.

### 1.2. Alternative Splicing Defects in Cancer Progression and Chemoresistance

Alternative splicing (AS) is an essential post-transcriptional process that involves the expansion of the cell’s protein-coding repertoire through the generation of multiple distinct mature mRNAs by a single gene *(Liu et al., 2017)*. Defects in splicing-regulatory elements and splicing factors, such as changes in expression levels or mutations, have been recently recognised as a hallmark of cancer *(Sveen et al., 2016)*. These defects as the primary contributors to therapeutic resistance in numerous cancer types are presently explored *(Duzgun and Oltean, 2025)*. Typical example includes the pyruvate kinase gene (PKM) splice variants that emerge under prolonged exposure to gemcitabine, resulting in the promotion of the PKM2 isoform through the selection of exon 10 *(Wang et al., 2012; Bai and Chen, 2020; Zahra et al., 2020)*.

Based on a multi-omics database for the case of pancreatic ductal adenocarcinoma, a high level of PKM2 isoform was found to be correlated with shorter recurrence-free survival and poor prognosis patients *(Calabretta et al., 2015)*. This splicing was subsequently shown to be guided by gemcitabine-induced upregulation of the polypyrimidine tract-binding protein (PTBP1), which serves as a major splicing factor that regulates PKM splicing, in chemoresistant pancreatic ductal adenocarcinoma cell lines. Reversing this splicing isoform towards the PKM1 isoform, which excludes exon 10, via knockdown of PTBP1, was able to restore sensitivity to gemcitabine in drug-resistant cells. Focusing on colon cancer, *Cheng et al. (2018)* also demonstrated enhanced sensitivity to vincristine and oxaliplatin in drug-resistant cancer cells through the regulation of glycolysis following the production of PKM1 isoform from silencing PTBP1 *(Cheng et al., 2018)*. These studies highlighted the promising prospects of PTBP1 as a therapeutic direction for drug-resistant cells resulting from splicing factor-induced AS, which has already been associated with patient clinical outcomes.

An integrative multi-omics analysis of clinical specimens prompts survival and differential analyses prior to experimental studies for the identification of novel potential targets linked to the development of drug resistance in cancer cells. This approach benefits future research on how to effectively overcome drug-resistance associated AS events in cancer cells.

### 1.3. The Role of SRPK1 in Chemoresistance

The discovery of the underlying causative roles and functional consequences of aberrant splicing is crucial to determine how splicing alterations contribute to drug resistance in cancer. Splicing decisions depend on various splicing factors phosphorylated by serine-arginine-protein kinases (SRPKs); hence, SRPKs have a critical role in regulating AS *(Das and Krainer, 2014; Patel, Sachidanandan, and Adnan, 2018)*.

Studies have linked **SRPK1**, a key regulator of AS, and its downstream targets to a wide spectrum of malignancies, including breast, colon, lung, prostate, and retinoblastoma *(Krishnakumar et al., 2008; Mavrou et al., 2014; Van Roosmalen et al., 2015; Gong et al., 2016; Yi et al., 2018)*. However, aberrant expression and functional alterations of SRPK1 appear to be related to both chemosensitivity and chemoresistance, raising an intriguing possibility on the role of SRPK1’s regulatory properties in stimulating the cellular response to anti-cancer drugs *(Nikas et al., 2019)*. *Odunsi et al. (2012)* observed restored sensitivity to cisplatin following the use of siRNAs to downregulate SRPK1, inhibiting ovarian cancer cell proliferation, migration, and invasion and promoting apoptosis. Conversely, *Schenk et al. (2001)* demonstrated reduced expression following the antisense oligonucleotide therapy targeting the translation initiation site of SRPK1, which enhanced ovarian cancer cell proliferation and cisplatin resistance *(Schenk et al., 2001)*. Hence, SRPK1 may be differentially modulated in distinct tumours; however, it remains largely unknown how tumour specific alterations contribute to chemoresistance mechanisms.

*Wang et al. (2020)* and *Huang et al. (2023)* revealed the vital role of SRPK1 in cancer, which, if derailed, may result in the development of tumour resistance. However, the expression landscape and clinical relevance of SRPK1 across different tumour types have not been systematically investigated. **In this article**, we aimed to explore multi-omics data from diverse tumour types to establish a robust foundation for a more comprehensive and clinically comparative analysis of SRPK1 alterations. This analysis provides an important direction for further understanding of the tumour specific mechanisms by which SRPK1 contributes to the development of chemoresistance, highlighting its multifaceted roles in this context.

## 2. Materials and Methods

### 2.1. Gene and Protein Expression Analysis

Using the “Exploration—Gene_DE” module of TIMER2 web (http://timer.comp-genomics.org/), SRPK1 expression in cancer was obtained from the boxplots of the expression level difference of SRPK1 between tumour and adjacent normal tissues for different tumours from the TCGA project. Meanwhile, the “Expression Analysis—Expression DIY—Box Plot” function of GEPIA2 *(Tang et al., 2017)* (Gene Expression Profiling Interactive Analysis, version 2) web tool (http://gepia2.cancer-pku.cn/#analysis), which produced boxplots of the expression level difference of SRPK1 between tumour and corresponding normal tissues from the GTEx database, was utilised to generate the controls in TIMER2 by matching breast invasive carcinoma (BRCA), colon adenocarcinoma (COAD) and prostate adenocarcinoma (PRAD) with normal tissues. For this, the cutoff values for *p*-value and log_2_FC (Fold Change) were set at 0.01 and 1, respectively.

Cancer proteomic datasets *(Chen et al., 2019)* from the UALCAN portal (http://ualcan.path.uab.edu/analysis-prot.html) were utilised for the analysis of the expression level of SRPK1 total protein between tumour and normal tissues in BRCA, COAD, and PRAD. In addition, the immunohistochemistry validation of differential protein expression of SRPK1 (Antibody No. HPA016431) between normal and tumour tissues in BRCA, COAD, and PRAD was conducted using the HPA database (https://www.proteinatlas.org/).

### 2.2. Survival Prognosis Analysis

The overall survival (OS) and disease-free survival (DFS) significance map data of SRPK1 in BRCA, COAD, and PRAD were obtained using the “Expression Analysis—Survival Analysis—Survival Map” function of GEPIA2. Based on the median (50% cutoff-high and 50% cutoff-low) of SRPK1 expression, the clinical patient cases of BRCA, COAD, and PRAD were divided into high– and low-expression group. Statistical significance was assessed through the log-rank (Mantel–Cox) test, which involved determining the hazard ratio (HR) with a 95% confidence interval (CI). Besides that, the “Expression Analysis—Survival Analysis” function of GEPIA2 was utilised to determine the Kaplan-Meier plots of OS and DFS in BRCA, COAD, and PRAD. Using a Kaplan-Meier Plotter tool (https://kmplot.com/analysis/index.php?p=background), the relationship between patient prognosis and SRPK1 expression in BRCA and COAD (PRAD was excluded in this case due to data unavailability) was examined based on the generated Kaplan-Meier survival curves for OS and relapse-free survival (RFS or also called DFS). Patients were grouped into high– and low-expression groups for survival analysis based on the “Auto select best cutoff” function.

### 2.3. SRPK1-Related Genes and Proteins Enrichment Analysis

Using the GeneMANIA *(Warde-Farley et al., 2010)* (https://genemania.org/) online portal, the function and association of SRPK1 were predicted for the construction of a gene–gene interaction network. The prediction output is in the form of a gene network that displays the top 20 interacting genes with SRPK1, where nodes symbolise genes and links represent networks.

A network analysis of SRPK1-binding proteins was subsequently searched on the STRING website (https://string-db.org/) *(Szklarczyk et al., 2015)* based on the following parameter settings: (1) the meaning of network edges was “confidence”; (2) the minimum required interaction score was “high confidence” (0.7); (3) the maximum number of interactions to show was “no more than 10 interactions” in the first shell. As a result, 10 proteins, including SRSF1, potentially interacted with SRPK1 were obtained. This study solely focused on SRSF1, a key regulator of alternative splicing, to evaluate its association with the survival information of patients with BRCA, COAD, and PRAD using the GEPIA2 and Kaplan-Meier Plotter databases, as described in Section 4.2.2.

Moreover, with the datasets of all TCGA tumours, the “Expression Analysis—Similar Gene Detection” function of GEPIA2 was utilised to assess the top 100 SRPK1-corrected targeting genes. Using the “Expression Analysis—Correlation Analysis” function of GEPIA2 in BRCA, COAD, and PRAD, Pearson’s correlation analysis of SRPK1 and the selected gene SRSF1 was then conducted. Through this module, dot plots were generated based on log2 [transcript per million (TPM)], yielding both *p-*values and correlation coefficients (R values).

### 2.4. Drug Sensitivity Analysis

The GSCA (https://guolab.wchscu.cn/GSCA/#/drug) pharmacogenomics platform *(Liu et al., 2018)*, which comprises the integration of gene expression profiles for 33 cancer types from the TCGA, and drug sensitivity data from the GDSC (Genomics of Drug Sensitivity in Cancer) dataset (the half-maximal inhibitory concentrations (IC_50_) of 251 small molecules in 1,001 cell lines) and CTRP (Cancer Therapeutics Response Portal) dataset (the area under the dose-response curve (AUC) values of 481 small molecules in 960 cell lines) datasets of various small molecules across different cell lines and corresponding SRPK1 mRNA levels were utilised to assess the correlation between drug sensitivity and SRPK1 mRNA expression levels. In both GDSC and CTRP datasets, sensitivity was classified as “negative” if higher SRPK1 expression correlated with lower IC_50_ or AUC values and “positive” if it was an inverse correlation. With a threshold of |r| ≥ 0.3 (i.e. being significantly positive correlation) or |r| ≤ −0.3 (i.e. being significantly negative correlation) and a false discovery rate (FDR) of ≤ 0.05, gene expression pattern in SRPK1 and drug sensitivity were correlated using the Pearson’s correlation coefficient.

Subsequently, the drug responses (log(IC_50_)) for patient samples in the TCGA cohorts based on their SRPK1 expression profiles were predicted using the CPADS (Comprehensive Pancancer Analysis of Drug Sensitivity) pharmacogenomics platform *(Li et al., 2024)*, which integrates gene expression profiles for 33 cancer types from the TCGA and corresponding drug sensitivity data from the GDSC dataset. As for the predicted drug responses, the “Drug Analysis—By Gene” function of CPADS was utilised to obtain predicted log(IC_50_) values for cisplatin in the BRCA and COAD patient cohorts and for docetaxel in the PRAD patient cohort. Not all agents would show clinical efficacy in the analysed cancer types; therefore, this study focused on cisplatin and docetaxel, as they are established as first-line treatments for these cancers.

Clinical patient samples of BRCA, COAD, and PRAD were grouped into high– and low-expression groups based on the median cutoff value of SRPK1 expression. For that, the two-sided Wilcoxon rank-sum test was conducted to evaluate the differences in median −log10 (IC_50_) values of each selected drug between high– and low-expression groups (based on the threshold of *p* < 0.05).

### 2.5. Statistical Analysis

The Wilcoxon rank-sum test was conducted to examine the mRNA expression levels of SRPK1 in tumour and normal tissues. Meanwhile, the SRPK1 protein levels between tumour and normal groups were compared using student’s *t*-test (two-tailed, unpaired). Besides that, log-rank (Mantel–Cox) test and Kaplan-Meier Plotter were employed for the prognostic relevance of SRPK1 expression. Additionally, the correlations of SRPK1 with various clinical features, such as SRSF1 and drug sensitivity, were examined through Pearson’s correlation analysis. Statistical significance was established at *p* < 0.05.

### 2.6. Ethical Approval and Consent to Participate

All data were obtained from publicly available databases that have already obtained ethical approval. Therefore, the analyses conducted in this study did not involve any ethical issues.

## 3. Results

An integrated multi-omics approach that took into account the unique SRPK1 expression profile of different tumour types and its drug sensitivity, functional interactions, and correlation with clinical prognostic value was utilised for this study’s pan-cancer analysis of SRPK1. **Figure 1** delineates the comprehensive framework of this multi-omics approach.

**Figure 1.**
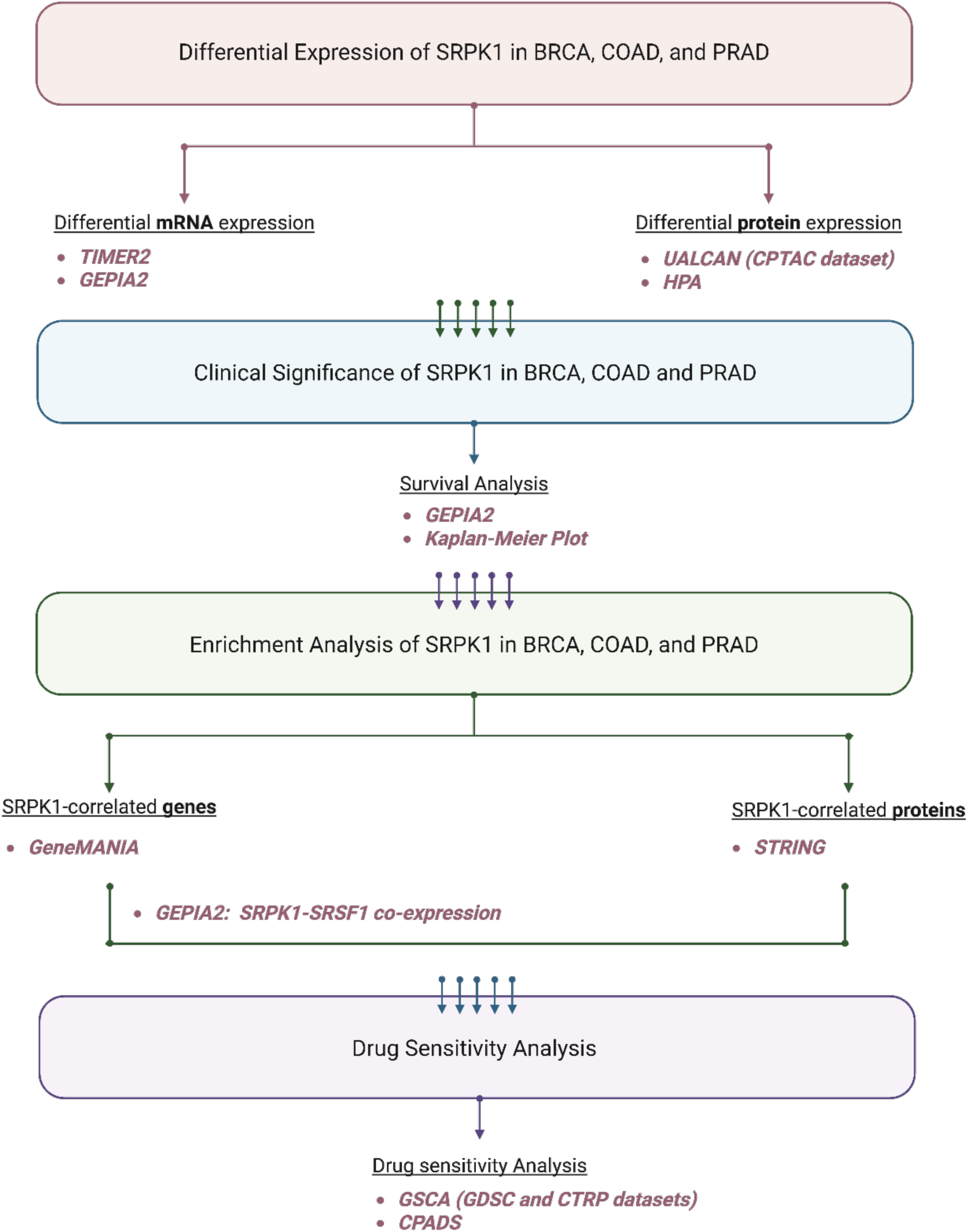
Summary of the study workflow and multi-omics databases used in SRPK1 analysis. This workflow illustrates the sequential steps of a comprehensive multi-omics analysis. The SRPK1 expression was first analysed at mRNA and protein levels, followed by the analysis on the relationship between survival probability and SRPK1 expression pattern, SRPK1-correlated genes and interacting proteins, and the impact of SRPK1 expression on drug sensitivity. Created with **<u>BioRender.com</u>**.

### 3.1. Tumour-Promoting Properties of Aberrant SRPK1 based on Transcriptomic and Proteomic Clinical Data in BRCA, COAD, and PRAD

Previous studies indicated that SRPK1 promotes tumorigenesis in various cancer types *(Malvi et al., 2020; Liu et al., 2021)*. The important question of whether aberrant expression of SRPK1 influences clinical outcomes in TCGA dataset remains unresolved. Thus, this study employed the pan-cancer analytical approach to examine the mRNA expression patterns of SRPK1 in cancer patients using TIMER2 database.

As shown in **Figure 2A**, as compared with normal tissues, SRPK1 mRNA expression was markedly upregulated in primary tissues for BLCA, BRCA, CESC, CHOL, COAD, ESCA, HNSC, LIHC, LUAD, LUSC, READ, STAD, and UCEC but downregulated in KICH and KIRC. However, no significant differences were detected in SRPK1 expression between primary tumours and paired normal tissues for GBM, KIRP, PAAD, PRAD, and THCA. Considering the frequent dysregulation of SRPK1 across various cancer types at transcriptomic level, the analysis then focused on its associations with specifically BRCA, COAD, and PRAD to better understand tumour-specific expression patterns underlying cancer heterogeneity. The limited number of normal tissue samples in TIMER2 database led to the analysis of SRPK1 mRNA expression in BRCA, COAD, and PRAD through the GEPIA2 platform, where normal tissues were incorporated with the GTEx dataset for more robust tumour-normal comparison. The analysis of GEPIA2 dataset revealed that BRCA and COAD achieved higher expression of SRPK1 in primary tumours, as compared with normal tissues, but this trend for PRAD achieved no statistical significance (*p* = 0.29; **Figure 2B**).

**Figure 2.**
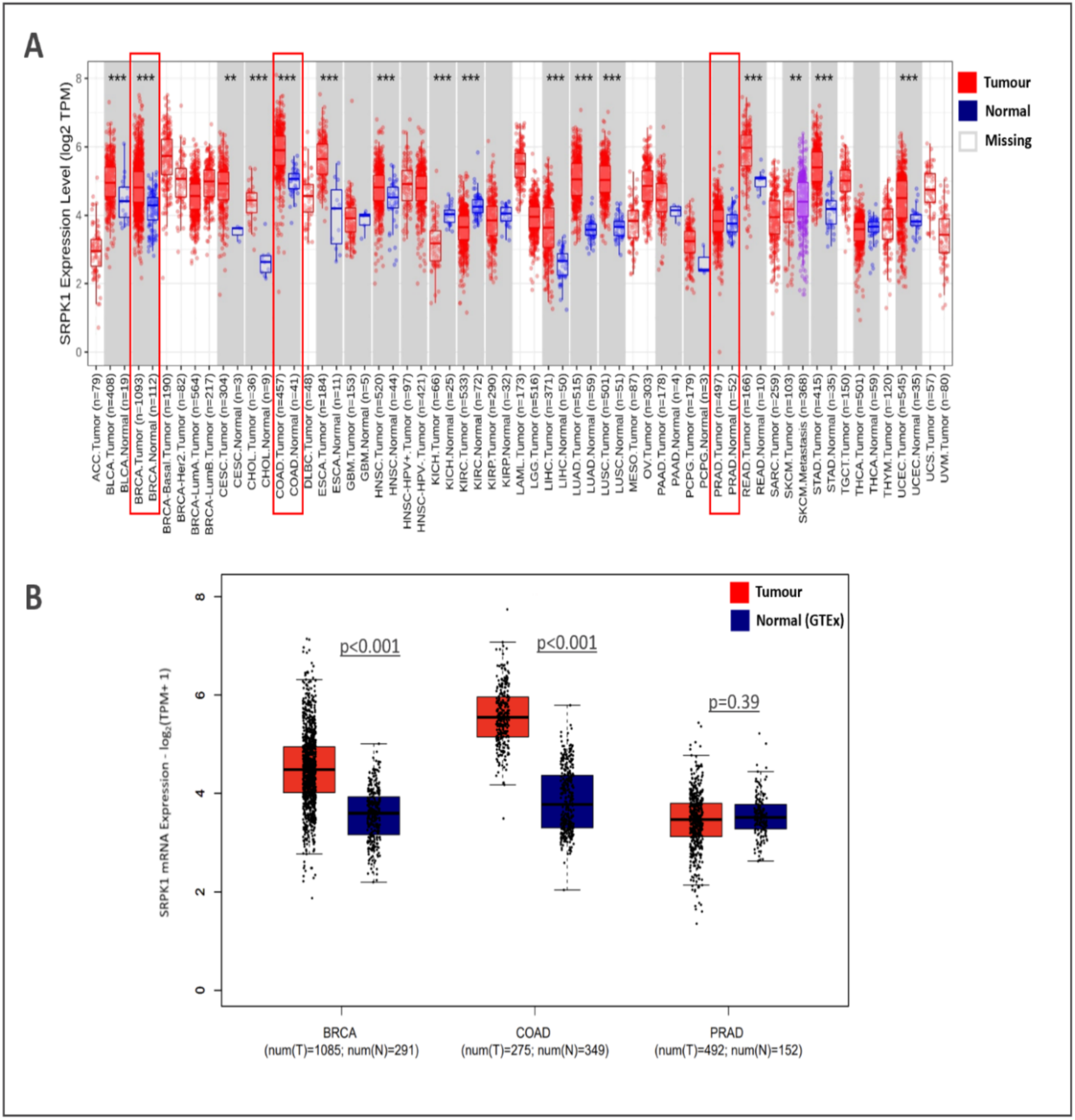
The differential mRNA expression of SRPK1 in pan-cancer samples. **(A)** SRPK1 mRNA levels in different **tumour** and corresponding **normal** tissues, obtained from TIMER2 database. SRPK1 expression levels are given on the y-axis, calculated as log2 (TPM; transcripts per million). The primary tumour and adjacent normal tissues are presented along the x-axis, together in a **grey** area. The Wilcoxon rank-sum test was conducted to determine statistical significance: **denotes *p* < 0.01; *** denotes *p* < 0.001. **(B)** Analysis of SRPK1 mRNA expression levels in primary tumour tissues of patients with BRCA [num(T) = 1085], COAD [num(T) = 275], and PRAD [num(T) = 492] in the TCGA dataset, as compared with those in normal-matched tissues for breast [num(N) = 291], colon [num(N) = 349], and prostate [num(N) = 152] in the GTEx dataset using the GEPIA2 platform. The transcriptional levels were log-normalised by the log_2_ (TPM+1) method. *P* < 0.05 was considered significant by descriptive statistics. These graphs use **red** boxplots to denote primary tumour tissues and **blue** box plots to denote adjacent normal tissues.

In addition to the transcriptomic alterations of SRPK1, protein expression CPTAC data quantified using the UALCAN platform were analysed. SRPK1 protein levels were compared between samples of primary tumour and paired adjacent normal tissues or age-stratified primary tumour tissues. In the case of primary tumour and adjacent normal tissues, the significantly higher SRPK1 protein levels in primary tumour samples in BRCA (*P* < 0.001; **Figure 3A**) and COAD (*P* < 0.001; **Figure 3B**) than their adjacent normal tissues were concordance with elevated mRNA levels for both BRCA and COAD. In the case of age-stratified primary tumour samples, PRAD patients aged 41–60 years and those aged 61–80 years recorded unchanged SRPK1 protein expression levels, possibly due to smaller sample sizes in loss of statistical power (*p* = 0.357; **Figure 3C**).

**Figure 3.**
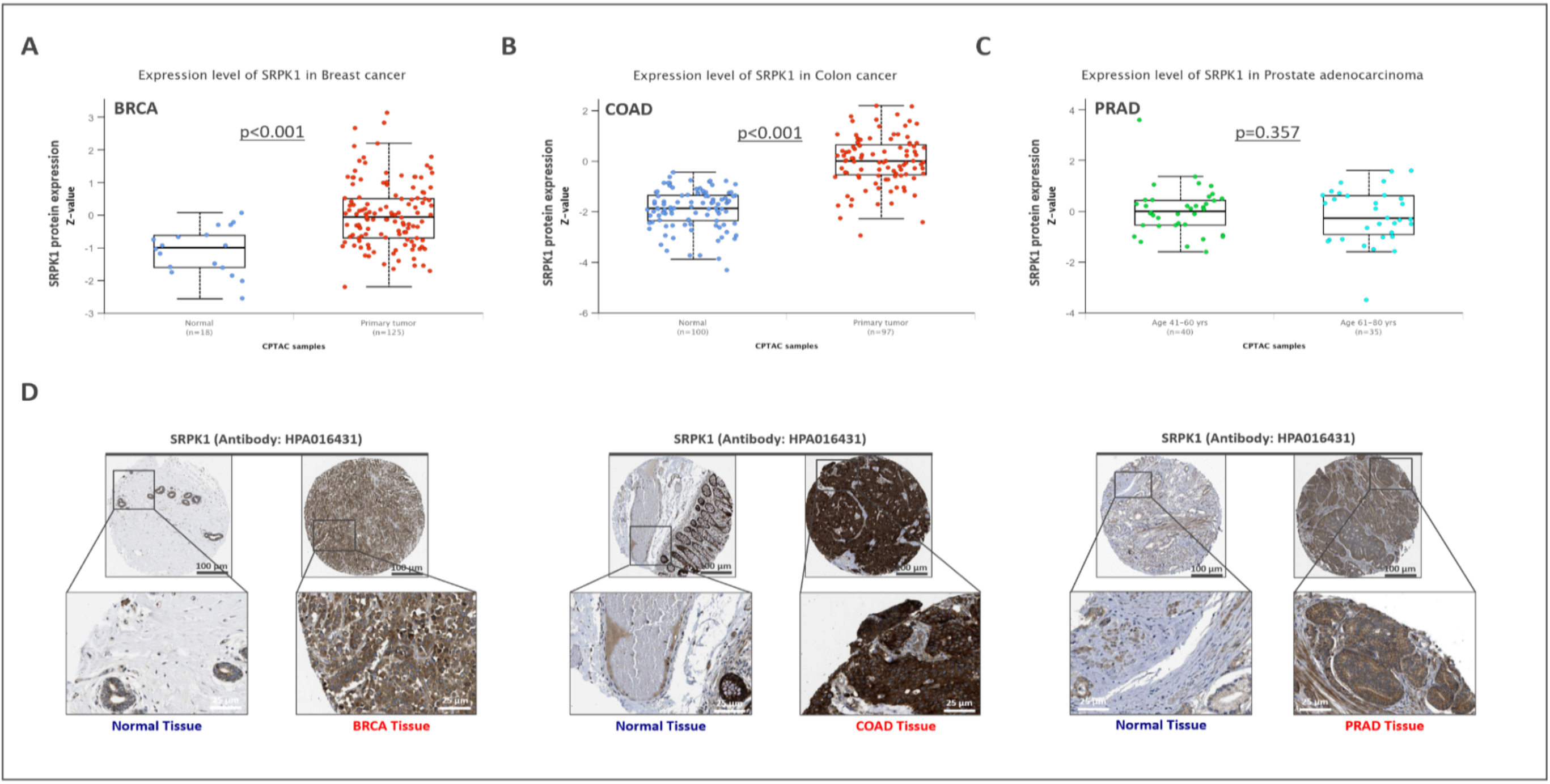
Differential expression analysis of SRPK1 protein levels in three different cancer tissues. (**A–B**) Comparison of SRPK1 protein expression between primary tumour tissues of BRCA (in **red**; n = 152) and COAD (in **red**; n = 97) and their corresponding normal breast (in **blue**; n = 18) and colon (in **blue**; n = 100) tissues from the CPTAC dataset on the UALCAN portal. **(C)** Analysis of the CPTAC-PRAD dataset for the protein expression of SRPK1 in age groups of 41–60 years (n = 40, highlighted in **green**) and 61–80 years (n = 35, highlighted in **light blue**). Z-values represent standard deviations from the median (the middle line in boxplot, with the lower and upper boundaries as the first and third quartiles and the whiskers extending to 1.5× the interquartile range of both quartiles) across samples for the given cancer types. Statistical significance (*p* < 0.05) of differential expression of SRPK1 was determined through an unpaired student’s *t*-test; *n* represents the number of samples in each boxplot. **(D)** Representative IHC images showing immunostaining for the SRPK1 protein in BRCA, COAD, and PRAD tissues and their corresponding normal tissues using HPA016431 antibody, obtained from the HPA dataset. Intense brown staining denotes increased expression in primary tumour tissues, as compared with normal tissues. Blue staining represents cell nuclei; brown staining shows SRPK1 protein deposits. Upper panel: scale bar = 100 µm; magnification 40×. Lower panel: scale bar = 25 µm; magnification 400×.

To further validate the protein levels of SRPK1 in those cancers, primary tumour tissues for which adjacent normal tissues were available were analysed using an alternative multi-omics HPA platform that included immunohistochemistry (IHC) staining data, where the degree of staining may more readily reflect protein levels. Based on the IHC staining of BRCA, COAD, and PRAD primary tissues for SRPK1, primary tumour tissues recorded significantly higher SRPK1 protein expression (mainly localised in the cytoplasm), as compared with the adjacent normal tissues (**Figure 3D**).

Taken together, clinical information and tissue samples were uneven across the many different pan-cancer types, but expression-based analysis revealed tumour-specific SRPK1 alterations and the potentially crucial role of SRPK1 in tumour progression. **Supplementary Table S1** lists the abbreviation of each cancer type.

### 3.2. Cross-Platform Identification of Gene-Expression-Defined Prognostic Features of SRPK1 in BRCA, COAD, and PRAD

Gene expression level associated to poor patient prognosis is indicative of that gene as a key contributor to the cancer progression *(Kaelin, 2017)* based on the assumption that, due to their conjectured role as cancer drivers in tumour-specific dependencies, prioritising gene expression changes associated with shorter survival times makes ideal therapeutic targets. In the following gene expression analysis, SRPK1 expression changes were identified as a significant adverse feature. Therefore, the analysis ensued by assessing whether these alterations could also be associated with poor prognosis in patient cohorts. GEPIA2 and Kaplan-Meier Plotter databases were utilised for the systematic analysis of the prognostic significance of SRPK1 expression status in BRCA, COAD, and PRAD cohorts. In the GEPIA2 database, the prognostic potential of SRPK1 was measured with a median cutoff point determined with HR for OS and DFS (in SRPK1-high-group versus SRPK1-low group for each cohort) (**Figure 4**).

**Figure 4.**
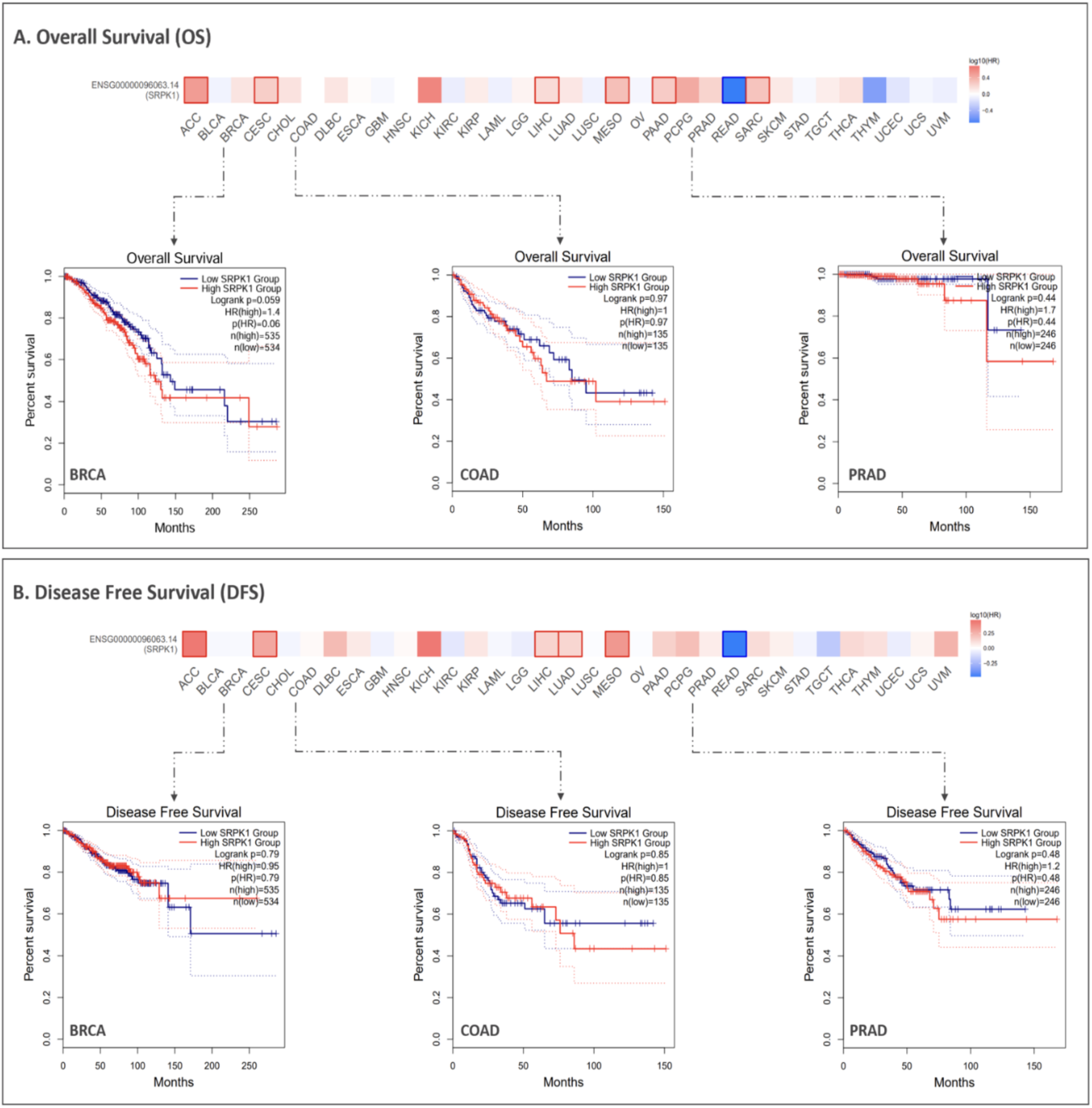
Survival analysis of patients with different SRPK1 mRNA expressions in BRCA, COAD, and PRAD clinical cohorts. **(A)** OS and **(B)** DFS for BRCA, COAD, and PRAD patients were analysed using the GEPIA2 database. Patients in the indicated cancer cohorts were stratified based on the median cutoff of SRPK1 mRNA expression, yielding Kaplan-Meier curves of OS and DFS (plotted in months). The **red** and **blue** lines represent the survival curves of patients with high and low levels of SRPK1 mRNA. Indicated by dashed lines, HR values (with 95% confidence intervals) represent the relative risk of the low-expression group compared with the high-expression group. HR > 1 implies the gene as a potential risk factor, whereas HR < 1 suggests a protective factor. Based on a log-rank (Mantel-Cox) test, a log-rank *p* < 0.05 indicates statistical significance. *n* indicates the patient count in each expression group.

Patients with low SRPK1 (n = 534) recorded a nearly significant longer OS than those with high SRPK1 (n = 535) in the BRCA dataset under the median cutoff point (HR = 1.4; log-rank *p* = 0.059; **Figure 4A, left**). In the COAD (HR = 1; log-rank *p* = 0.97; **Figure 4A, middle**) and PRAD (HR = 1.7; log-rank *p* = 0.44; **Figure 4A, right**) datasets, with 135 and 246 patients respectively, a better OS was observed in patients with high SRPK1 than those with low SRPK1 despite the statistically insignificant differences.

Following that, a DFS analysis of all three cohorts was conducted, which revealed a trend towards high expression of SRPK1 being associated with disease progression in the BRCA dataset (HR = 0.95; log-rank *p* = 0.79; **Figure 4B, left**). As for the COAD (HR = 1; log-rank *p* = 0.85; **Figure 4B, middle**) and PRAD (HR = 1.2; log-rank *p* = 0.48; **Figure 4B, right**) datasets, high expression of SRPK1 was not associated with adverse outcomes. However, these associations were not statistically significant, which may be attributed to the small sample size of patients in clinical settings.

Therefore, this study considered the possibility that the utilisation of other related databases with larger datasets associated with BRCA, COAD, and PRAD patient survival outcomes may uncover prognostic relationships not found in this study’s GEPIA2 analysis. Addressing that, this study proceeded with an additional prognostic analysis using the Kaplan-Meier Plotter database, which contains larger patient datasets, to evaluate survival-associated gene expression patterns at the mRNA level. For Kaplan-Meier OS and DFS analyses, an optimal clinically relevant cutoff was applied. Patients were dichotomised into SRPK1-high and –low groups in the BRCA and COAD datasets (**Figure 5**) based on maximally selected log-rank statistics yielding hazard ratios. Please note that, in this database, corresponding data were not available for PRAD.

**Figure 5.**
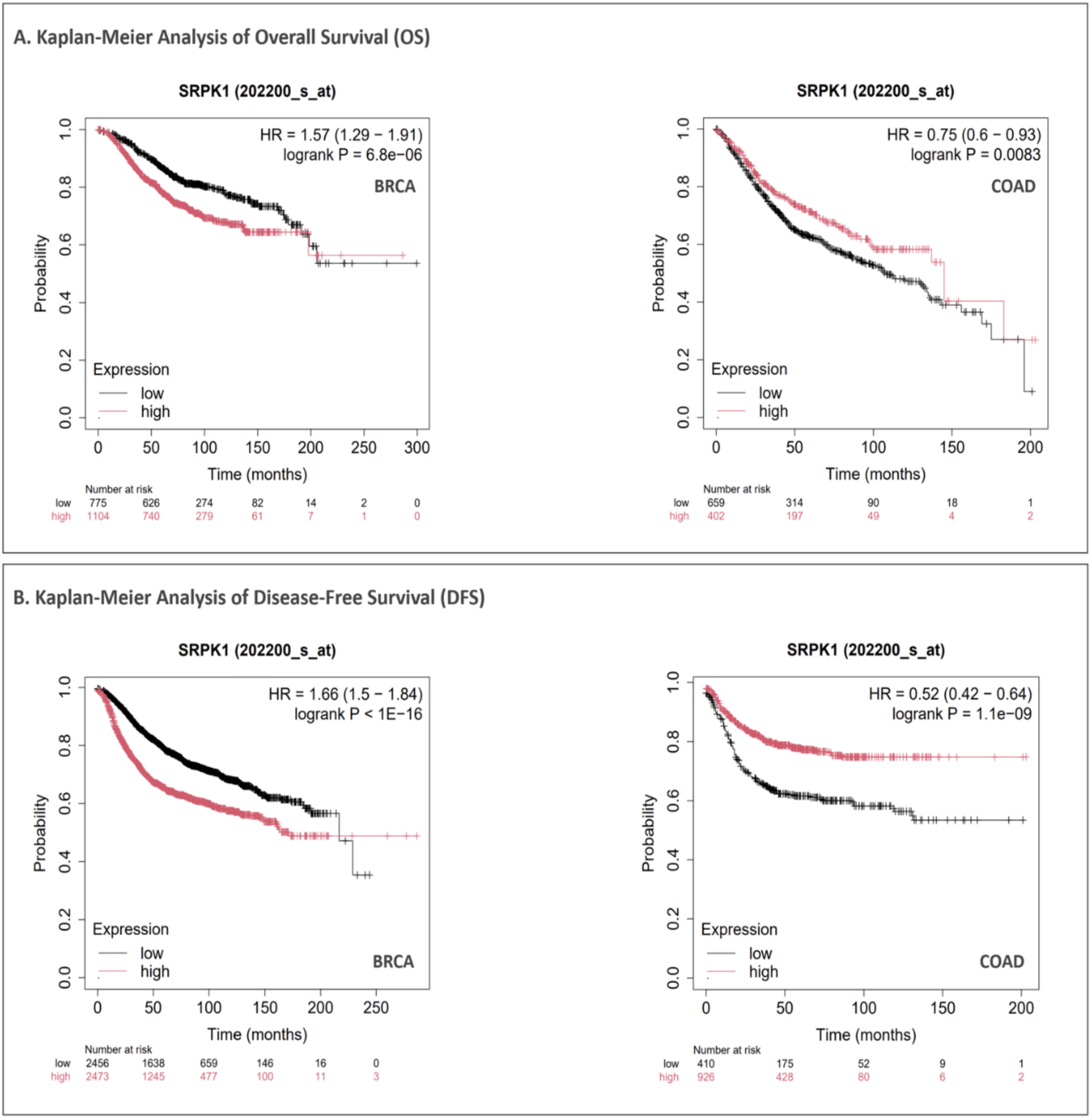
Analysis of the expression and prognosis relevance of SRPK1 in BRCA and COAD cohorts. Kaplan-Meier plots revealed differences in **(A)** OS and **(B)** DFS of BRCA and COAD patients based on the 202200_s_at probe obtained from the Kaplan-Meier plotter database. The OS and DFS of BRCA and COAD patients stratified by low (**black** line) and high (**red** line) SRPK1 expressions using the optimal cutoff point provided by Kaplan-Meier plotter database. The OS and DFS of high– and low-expression groups were compared based on the HR (with 95% confidence intervals). Log-rank (Mantel–Cox) test was performed to determine statistical significance; *p* < 0.05 indicates statistical significance.

Results of Kaplan-Meier survival analysis revealed significantly worse OS in BRCA patients with high SRPK1 expression (n = 1104) than those with low SRPK1 expression (n = 775) (HR = 1.57; log-rank *p* = 6.8e-06; **Figure 5A, left**). A similar trend was observed for DFS (high SRPK1: n = 2473, low SRPK1: n = 2456; HR = 1.66; log-rank *p* < 1E-16; **Figure 5B, left**). This suggested the potential pro-tumour role of upregulated SRPK1 in breast carcinogenesis. Meanwhile, low SRPK1 expression was found to be correlated with a marked decrease in OS (high SRPK1: n = 402, low SRPK1: n = 659; HR = 0.75; log-rank *p* = 0.0083; **Figure 5A, right**) and DFS (high SRPK1: n = 926, low SRPK1: n = 410; HR = 0.52; log-rank *p* = 1.1e-09; **Figure 5B, right**) in patients with COAD. This indicated that downregulation of SRPK1 may contribute to the progression of colon cancer. Collectively, this disparity reflected the complex and multifaceted role of SRPK1 expression in mediating cancer-type-dependent associations between clinical variables and poor outcomes.

### 3.3. Identification of Gene–Protein Interactions Associated with SRPK1-Bound Functional Networks Underpinning Alternative Splicing

Cancer cells exhibit extensive heterogeneity in their cell states, characterised by altered gene and protein expression signatures associated with epithelial-to-mesenchymal transition (EMT), invasion, metastasis, aberrant RNA splicing, and therapeutic resistance, among other oncogenic processes *(Marusyk, Janiszewska, and Polyak, 2020; Patel and Yanai, 2024)*. This study’s preceding integrative multi-omics clinical data analyses suggested that the abnormal expression of SRPK1 mRNA and protein is broadly associated with unfavourable outcomes in cancers. This prompted this study to explore the functional network of genes and proteins interacting with SRPK1, specifically on whether the identification of gene–gene and protein–protein interaction networks could provide deeper insights on the SRPK1-dependent biological functions.

In order to identify gene(s) closely related to the regulatory functional network in which SRPK1 was involved, this study first performed a gene–gene interaction analysis using the GeneMANIA database on the top 20 genes most correlated with SRPK1. These genes were categorised as pan-cancer genes and exhibited a number of gene–gene interactions, which formed a densely interconnected ‘‘main’’ physical network, reflecting potential functional associations among SRPK1-bound genes shown in **Figure 6A**. Functional gene–gene interaction analysis across pan-cancer tumour samples revealed 20 SRPK1-related genes enriched in numerous cellular processes, including mRNA splicing and processing (*ZRSR2*, *RBM4*, *SRSF1*, *SF3A3*, *SRSF6*, *PRPF8*, *PRPF38A*, *TARDBP*), chromatin organisation and transcriptional regulation (*LBR*, *SAFB*, *SAFB2*, *NAT10*, *SIRT7*), ubiquitin-mediated proteolysis and protein homeostasis (*HECW2*, *PPM1G*), DNA damage response and cell cycle regulation (*GTF3C4*, *GBP2*), chromatin remodelling in germ cell development (*PRM1*), and immune response and signal transduction (*GBP2*, *SERPINB10*). These processes are crucial in therapeutic resistance *(Sheng et al., 2018; Pan et al., 2023; Xie et al., 2023)*, tumour development, and progression *(Ren et al., 2022; López-Cánovas et al., 2023; Yi et al., 2023)*.

**Figure 6.**
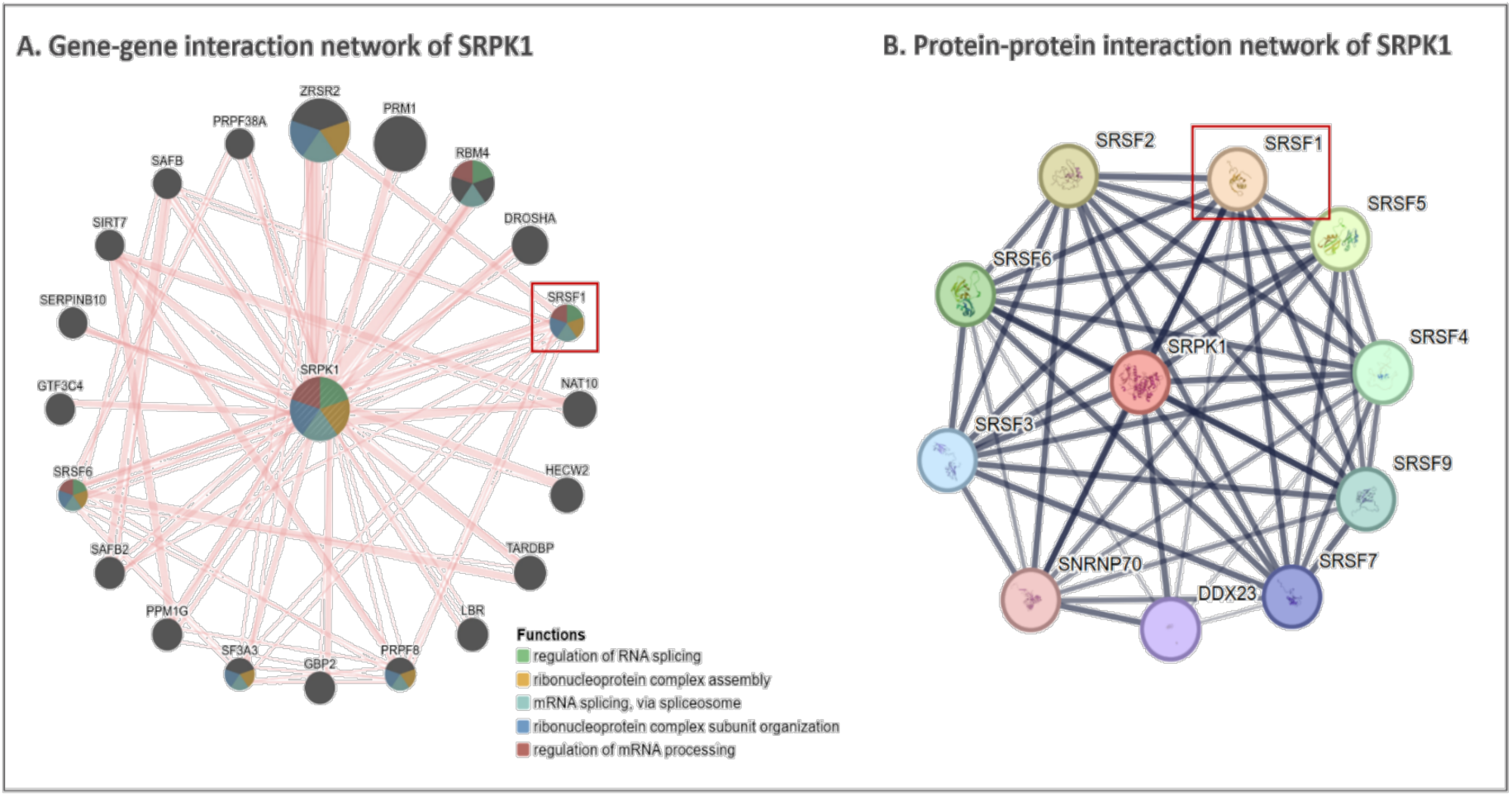
Functional enrichment analysis of SRPK1-interacting genes and proteins across pan-cancers. **(A)** Gene–gene interaction network of SRPK1 obtained from the GeneMANIA database illustrates its physical and functional partners. In the network, genes are represented by **dark-grey** nodes, and their interactions are denoted by **pink** lines between nodes (edges). A node is represented by a pie chart that indicates the distribution of each gene’s functional categories associated with SRPK1. Pie chart colours: **green** denotes regulation of RNA splicing; **yellow** denotes ribonucleoprotein complex assembly; **aqua** denotes mRNA splicing via spliceosome; **blue** denotes ribonucleoprotein complex subunit organisation; **red** denotes regulation of mRNA processing. The selected gene, SRSF1, which exhibited the strongest functional association with SRPK1 among the interacting genes, is framed in a **red** square. Colour coding of the pie chart pieces is shown on the right side of the figure. **(B)** Protein–protein interaction network of SRPK1, generated using the STRING physical interactions database, identifies functional interactions associated with the pre-mRNA splicing process in tumours. Each node (coloured circle) represents a group of SRPK1-interaction proteins with functional relationships, and a **black** edge indicates an interaction. Node colours do not have any meaning. Protein–protein interactions were identified at a high confidence score (≥ 0.7). The thickness of the edges reflects the strength of supporting evidence, where thicker edges indicate higher interaction scores. The selected protein, SRSF1, outlined in a **red** rectangle, represents the strongest predicted interaction.

Next, a protein–protein interaction analysis was conducted using the STRING database based solely on experimentally validated sources to identify key proteins functionally interacting with SRPK1 across different cancer types (**Figure 6B**). This analysis revealed that a network consisting of serine/arginine-rich (SR) splicing factors, SRSF1, SRSF5, SRSF4, SRSF9, SRSF7, SRSF3, SRSF6, and SRSF2, the spliceosomal RNA helicase DDX23, and the U1 snRNP component SNRNP70, which strongly interacted with SRPK1, may mediate drug response by regulating the alternative splicing network. E2F1 transcriptionally activated DDX23, a splicing factor, which regulated the mRNA processing of FOXM1, promoting the progression of ovarian cancer progression *(Zhao et al., 2021)*. The overexpression of SNRNP70, a nuclear RNA-binding protein regulating splice variants of transcripts, may drive the formation and growth of osteosarcoma by regulating the skipping of exon 10 of CD55 (*Li et al.*, *2024*). Among them, serine/arginine-rich (SR) family members—well-established drivers of tumorigenesis—particularly SRSF1, were highlighted, and their abnormal expression in various cancer types pointed to chemoresistance or drug-tolerance mechanisms by producing impaired alternative splicing patterns *(Sheng et al., 2018; Pellarin et al., 2020; Sinnakannu et al., 2020; Zhang et al., 2021; Wang et al., 2022; Bian et al., 2024)*.

Taken together, analysing these databases revealed that, beyond the growing recognition of the importance of abnormal RNA splicing in tumorigenesis, the identification of its gene–gene and protein–protein interaction networks captures a functional insight, revealing the potential role of SRPK1 as a critical contributor to therapeutic resistance. The description of gene functions is listed in **Supplementary Table S2**.

### 3.4. Co-Expression of the Selected Splicing Factor SRSF1 with SRPK1 and Association with its Prognostic Significance in BRCA, COAD, and PRAD

Previous studies demonstrated that SRPK1 and SRPK1-associated splicing regulators are often aberrantly expressed in cancers and can drastically affect cancer malignancy through both the splicing-regulatory network and, at least in part, effective co-expression interactions with one another *(Wagner et al., 2019; Liu et al., 2021)*. There are sporadic findings describing alterations in the expression and corresponding functions of these splicing regulatory elements in different malignancies. Nevertheless, clinically relevant data on their gene-level correlations in the context of their overexpression or downregulation should assist in dissecting their cancer-specific actions.

As the first step towards this, this study focused on splicing factor SRSF1, a prototypical SR family member, among the identified SRPK1-related genes/proteins (**Figure 7**). Apart from being broadly expressed in distinct cell types, it regulates the splicing pattern of many important genes. As for the dissection of the SRPK1-SRSF1 interaction at the mRNA level, their co-expression patterns in BRCA, COAD, and PRAD patient cohorts were analysed using the GEPIA2 database. Correlation analysis for COAD (*R* = 0.63; *p* < 0.001; **Figure 7B**) and PRAD (*R* = 0.68; *p* < 0.001; **Figure 7C**) patient samples revealed a strong positive correlation between SRPK1 and SRSF1 mRNA expression levels. The same results were obtained in BRCA patient samples (*R* = 0.45; *P* < 0.001; **Figure 7A**), but its correlation was moderate, as compared with that of COAD and PRAD patient samples. Results suggested the potential influence of SRSF1 on SRPK1-driven AS via a combination of mechanisms when co-expressed with SRPK1 in the analysed cancer types.

**Figure 7.**
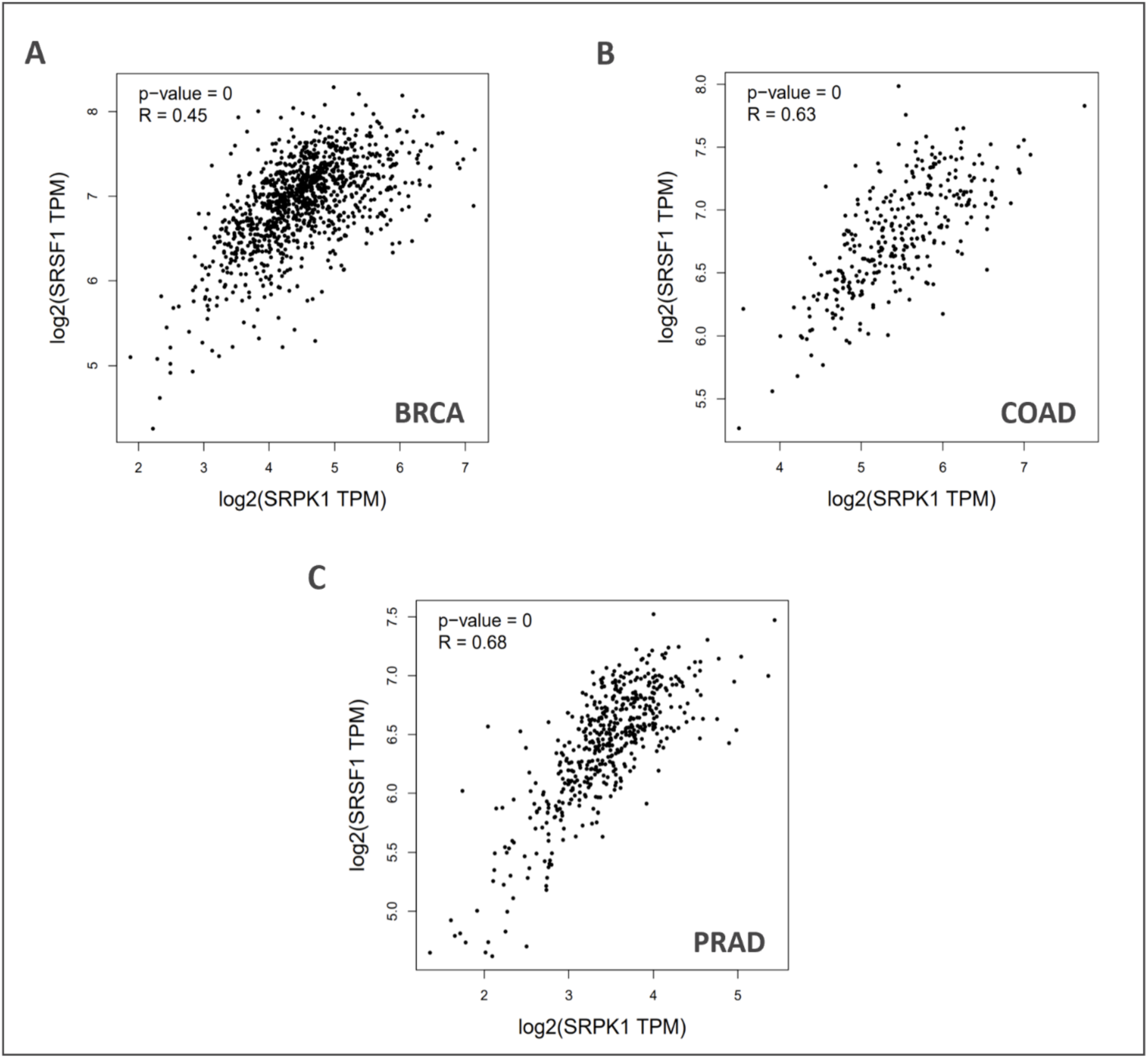
Selected splicing factor SRSF1 positively correlated with SRPK1 in BRCA, COAD, and PRAD patient samples. Scatterplots exhibit the correlation between SRPK1 (x-axis) and SRSF1 (y-axis) expression at the mRNA level based on **(A)** TCGA-BRCA, **(B)** TCGA-COAD, and **(C)** TCGA-PRAD datasets analysed using the GEPIA2 database. The correlation coefficient (R value) was calculated using Pearson’s correlation, and the corresponding *p-*value indicates the statistical significance of the correlation. Each **black** dot represents the log2-transformed expression levels (TPM; transcripts per million) of both genes in an individual patient.

Meanwhile, the survival analysis of SRPK1 revealed the expression level of SRPK1, with significant association with pour outcome in cancer-type specific survival, as the most prominent prognostic feature. Given its predominant positive correlation with SRSF1, this study proceeded to examining the prognostic significance of SRSF1 expression in BRCA, COAD, and PRAD patient cohorts using two independent databases, namely GEPIA2 and Kaplan-Meier Plotter databases.

This study first used the GEPIA2 database to assess the alterations in SRSF1 mRNA expression in relation to OS and DFS for BRCA, COAD, and PRAD patients, where tumour samples were categorised into high– and low-expression subgroups based on the median SRSF1 expression level in each patient cohort (as the cutoff value) (**Supplementary Figure S1**). The survival analysis demonstrated that high levels of SRSF1 tended to coincide with shorter OS in both BRCA (HR = 1.1; log-rank *p* = 0.49; **Supplementary Figure S1A, left**) and PRAD (HR = 1.8; log-rank *p* = 0.37; **Supplementary Figure S1A, right**) patient cohorts (n = 1,070 and n = 492, respectively) despite the statistical insignificance. In contrast, the data of the COAD patient cohort yielded a substantially inferior OS (HR = 0.55; log-rank *p* = 0.015; **Supplementary Figure S1A, middle**) for patients with SRSF1-low expression tumours (n = 135) compared with what was seen in those with SRSF1-high expression tumours (n = 135) although the number of tumour samples in both expression subgroups was rather low. Moreover, based on the results of DFS analyses, PRAD patients with SRSF1-low expression tumours tended to have better survival outcomes (n = 492; HR = 1; log-rank *p* = 0.83; **Supplementary Figure S1B, right**), whereas BRCA patients with SRSF1-low expression tumours tended to have worse clinical survival outcomes (n = 1,070; HR = 0.89; log-rank *p* = 0.55; **Supplementary Figure S1B, left**). A similar pattern to the BRCA patient cohort was observed in the COAD patient cohort (n = 270; HR = 0.72; log-rank *p* = 0.17; **Supplementary Figure S1B, middle**).

Overall, the difference in DFS between SRSF1-high expression group and SRSF1-low expression tumour group was not statistically significant. When analysing separately within the BRCA, COAD, and PRAD cohorts, most of the associations became insignificant, likely due to the relatively sample size in each expression subgroup.

However, the small sample size involved, especially in COAD and PRAD patient cohorts, may obscure the prognostic effects associated with SRSF1 expression levels in the above analysis. With that, this study examined the associations of SRSF1 mRNA expression levels with OS and DFS by leveraging the large patient cohorts whose survival times were available in the Kaplan-Meier Plotter database. Tumour samples from BRCA and COAD cohorts were divided into high– and low-expression subgroups according to the optimal cutoff point. However, OS and DFS could not be assessed due to the unavailability of data for the PRAD cohort in the Kaplan-Meier Plotter database (**Supplementary Figure S2**).

In this analysis, a trend towards high levels of SRSF1 expression in tumour samples was found to be associated with better OS in both BRCA (n = 943; HR = 0.77; log-rank *p* = 0.056; **Supplementary Figure S2A, left**) and COAD (n = 814; HR = 0.84; log-rank *p* = 0.16; **Supplementary Figure S2A, right**) patient cohorts despite the statistical insignificance. Kaplan-Meier survival analysis further demonstrated that both BRCA (n = 2032; HR = 0.64; log-rank *p* = 5.5e-09; **Supplementary Figure S2B, left**) and COAD (n = 1167; HR = 0.66; log-rank *p* = 0.00045; **Supplementary Figure S2B, right**) cohorts exhibited significantly longer DFS in patients with high-SRSF1 expression, as compared with those with low-SRSF1 expression. These results suggested that changes in SRSF1 expression levels may be at least partially associated with poor patient outcomes, as a crucial fraction of this association reflected a cancer-type-specific manner, thereby highlighting the heterogeneity across cancer types.

Taken together, these results suggested that SRPK1-driven expression of SRSF1 (i.e., their co-expression) may act in a promalignant fashion for the analysed cancer types. As abnormal expression of splicing factors are known to promote the likelihood of either exon skipping or inclusion in AS, this cooperation potentially leads to even higher levels of an oncogenic isoform; thus promoting tumour maintenance and treatment resistance-associated processes across multiple cancer types. **Supplementary Figure S3** presents the schematic representation of the proposed model of action of the SRPK1-SRSF1 co-expression axis.

### 3.5. Pharmacogenomics Insights on SRPK1 Expression-Driven Sensitivity and Resistance to Anti-Cancer Agents

It has long been suggested that cancer-specific responses to anti-cancer agents may be dependent on changes in gene expression *(Roy et al., 2019; Shea et al., 2025)*. Gene expression changes in splicing regulatory elements and/or splicing factors, both individually and in coordination, have been shown to contribute to the frequent changes in the splicing patterns of target genes across cancers and cell types, which may ultimately lead to divergent responses to anti-cancer agents *(Deng et al., 2021; Anczukow et al., 2024)*. Findings on the alteration of SRPK1 in a cancer-specific manner and its association with poor patient prognosis and expression of distinct splicing isoforms prompted this study to assess whether changes in SRPK1 expression levels may be among the unknown driving forces behind the drug-resistance mechanisms. Therefore, this study utilised independent pharmacogenomics datasets to determine the extent of and how the relationships between changes in SRPK1 expression and responses to various anti-cancer agents in both **cancer cell lines** and **patients’ tumour samples**.

Based on the independent drug sensitivity datasets from the GDSC and CTRP, a systematic analysis was conducted to assess the influence of SRPK1 expression on the response of individual cell lines to targeted agents. This analysis included 1,001 cancer cell lines and 251 unique anti-cancer agents from the GDSC dataset and 960 cancer cell lines and 481 unique anti-cancer agents from the CTRP dataset. Cancer cell lines in both datasets are of various lineages, including breast, ovarian, colon, skin, blood, and prostate. Additionally, the anti-cancer agents target a broad range of cancer-related processes, including phosphonisitide 3-kinase (PI3K)/mammalian target of rapamycin (mTOR) signalling, extracellular signal-regulated kinase 1 (ERK)/mitogen activated protein kinase (MAPK) signalling, receptor tyrosine kinase (RTK) signalling, epidermal growth factor receptor (EGFR) signalling, apoptosis pathway, EMT, DNA damage response, and cell cycle.

Given the various tissue-specific lineages and the diversity of anti-cancer agent targets available in either dataset, an evidence-based approach was adopted, in which drug sensitivity data provides meaningful insights on the gene expression-based dependencies of drug responses. The effect of each anti-cancer agent on gene expression-based cell line sensitivity was assessed based on the IC_50_ value (drug concentration at 50% viability) from the GDSC dataset and the AUC value (area under the dose-response curve) from the CTRP dataset (where IC50 values were not reported).

As shown in **Figure 8A**, the analysis of the GDSC dataset for hundreds of cancer cell lines revealed the association of SRPK1-high levels with either sensitivity or resistance to the top 30 important anti-cancer agents (out of 251 tested anti-cancer agents). Of these, nine anti-cancer agents were positively correlated with SRPK1-high levels, suggesting that overexpression of SRPK1 may render multiple cancer cell lines more resistant to these agents. Conversely, 21 anti-cancer agents were negatively correlated with SRPK1-high levels, indicating that increased SRPK1 levels may enhance the sensitivity of diverse cancer cell lines to these agents. However, the relatively weak correlations between SRPK1 overexpression and alterations in response to all 30 anti-cancer agents (Pearson’s correlation coefficient *r* ranging from –0.25 to 0.25; **Supplementary Table S3**) were observed despite being statistically significant (FDR ≤ 0.05). Notably, in cancer-specific cell lines, SRPK1 expression levels were most strongly and positively correlated with resistance to TGX221 (*r* = 0.243; FDR = 2.05E-05), followed by erlotinib (*r* = 0.230; FDR = 0.000246) and lapatinib (*r* = 0.214; FDR = 0.000207), and most strongly and negatively correlated with sensitivity to NPK76-II-72-1 (*r* = −0.266; FDR = 1.87E-15), PHA-793887 (*r* = −0.227; FDR = 1.9E-11), and AT-7519 (*r* = −0.209; FDR = 9.41E-10). Nonetheless, several observed anti-cancer agents have been reported to exhibit treatment responses associated with changes in gene expression levels, such as erlotinib *(Yamasaki et al., 2007; Lv et al., 2019; Pal et al., 2021)* and lapatinib *(Goel et al., 2016; Ruprecht et al., 2017; Jiang et al., 2018; Steggall et al., 2025)*, as well as novel anti-cancer agents whose associations are yet to be clearly identified, including the top-ranked PHA-793887 and AT-7519. However, none of these anti-cancer agents have been linked to SRPK1.

**Figure 8.**
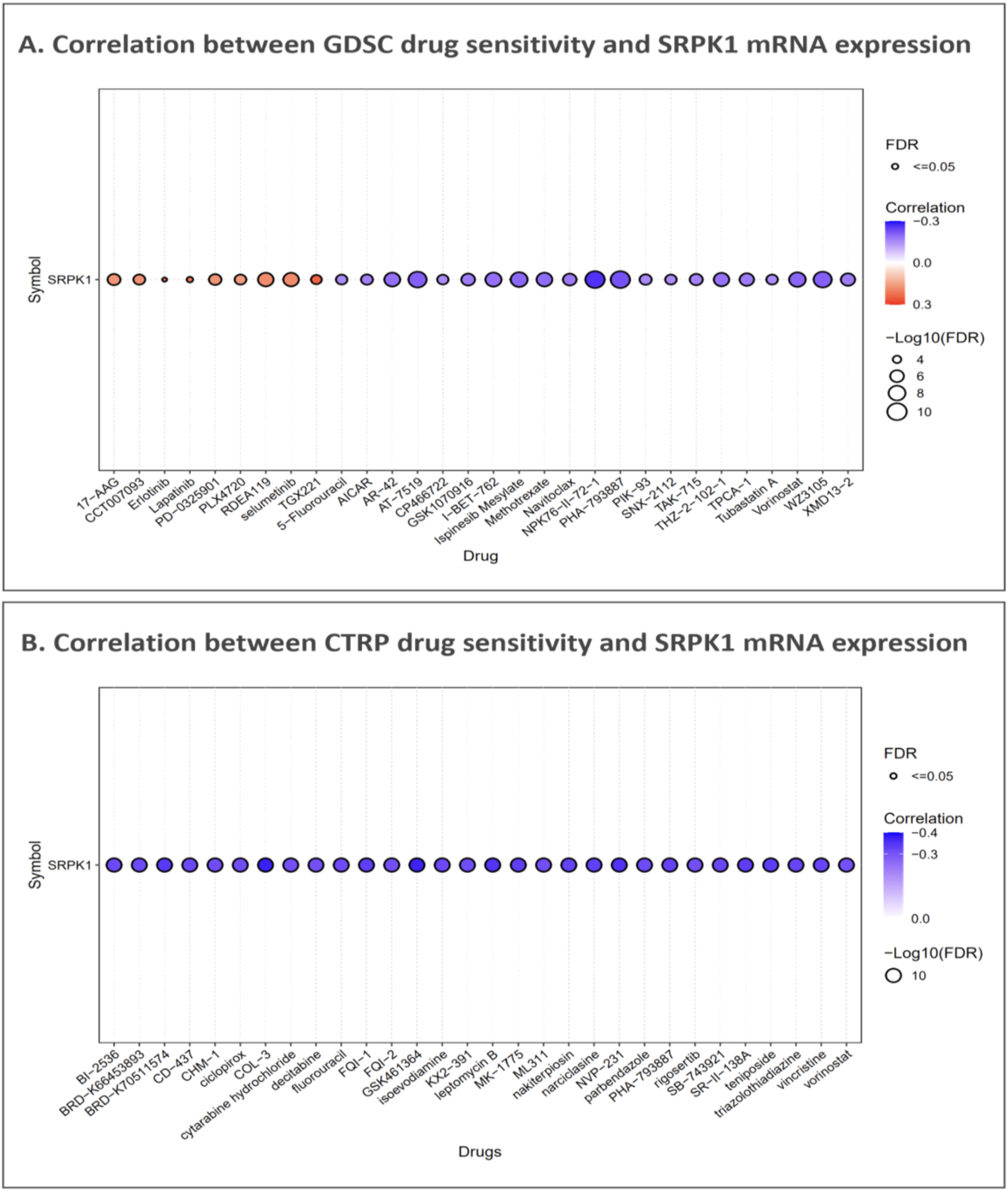
SRPK1 mRNA expression in relation to anti-cancer drug sensitivity across multiple cancer cell lines based on two independent pharmacogenomics datasets. **(A)** Bubble plot representation of the top 30 agents correlated (either positively or negatively) with SRPK1 mRNA expression across 1,001 cancer cell lines in the GDSC drug sensitivity dataset, comprising a total of 251 anti-cancer agents. **(B)** Bubble plot depicting the negative correlations of the top 30 agents associated with SRPK1 mRNA expression across 960 cancer cell lines in the CTRP drug sensitivity dataset, comprising a total of 481 anti-cancer agents. The correlation coefficients were calculated using Pearson’s correlation. The corresponding FDR values indicate the statistical significance of the correlations. The gradient colour bar shows the correlation coefficient; bubble size indicates the significance of the correlation, reflected by −log_10_ FDR. The bubble colour intensity signifies the degree of association with anti-cancer agent sensitivity and resistance. Red bubbles indicate a positive correlation (i.e., increased resistance to the specified agent in cancer cell lines with higher SRPK1 expression), and blue bubbles represent a negative correlation (i.e., increased sensitivity to the specified agent in cancer cell lines with higher SRPK1 expression).

Decreased H19 expression has been shown to upregulate vimentin and downregulate E-cadherin concurrently, leading to EMT and subsequent resistance to the EGFR inhibitor erlotinib in the EGFR-mutated non-small-cell lung cancer (NSCLC) cell line, HCC827 *(Chen et al., 2020)*. In another NSCLC cell line, A549, erlotinib-induced resistance has been associated with H19-high levels, which promote the expression of ATG7 by inhibiting miR-615-3p, thereby increasing the likelihood of autophagy activation *(Pan and Zhou, 2020)*. Together, these studies suggested that tissue– and lineage-specific changes in gene expression are likely to modulate the sensitivity or resistance to anti-cancer agents. Considering the lineage-specific findings mentioned above, this may explain why no strong correlations between sensitivity/resistance and SRPK1 expression levels were observed. A possible explanation for our GDSC observation could be that pan-cancer associations, with a large and heterogeneous collection of cell lines, dilute out the strength of cancer-specific correlations that might arise due to lineage-specific differences in SRPK1 expression levels. This indicates that sensitivity and resistance to many anti-cancer agents may be modulated by expression changes within the context of cancer-type-specific lineages. Subsequently, analysis of the CTRP dataset for cancer cell lines revealed that SRPK1-high levels were associated with sensitivity to the top 30 important anti-cancer agents (out of 481 tested anti-cancer agents), as shown in **Figure 8B**. We examined this in more detail; all 30 anti-cancer agents showed negative correlations with SRPK1-high levels, indicating that increased SRPK1 levels may enhance the sensitivity of diverse cancer cell lines to these agents. In addition, significant (FDR ≤ 0.05) yet moderate correlations between SRPK1 overexpression and alterations in response to all 30 anti-cancer agents (Pearson’s correlation coefficient *r* ranging from – 0.31 to 0.38; **Supplementary Table S4**) were observed.

The majority of these associations are novel observations derived from the CTRP dataset; in other words, none of these anti-cancer agents have previously been linked to SRPK1. Altogether, this suggests that increased SRPK1 expression might enhance drug sensitivity across diverse cancer cell lines.

To understand whether differences in drug response are due to gene expression changes, we partitioned TCGA samples from the BRCA, COAD, and PRAD patient cohorts into SRPK1-high and SRPK1-low groups using the CPADS platform and then assessed the differences in predicted IC50 values between the two groups in the analysed cancer types (**Figure 9**). We examined SRPK1 expression and its relationship with predicted cisplatin IC_50_ values in BRCA and COAD cohorts, as well as with predicted docetaxel IC_50_ value in PRAD cohort. Notably, COAD patients in the SRPK1-low group exhibited significantly higher cisplatin IC_50_ values compared to the SRPK1-high group; therefore, decreased SRPK1 expression may contribute to cisplatin resistance in COAD patients (*p* = 0.0065; **Figure 9, middle**). In contrast, PRAD patients in the SRPK1-high group showed significantly higher docetaxel IC_50_ values, suggesting that overexpression of SRPK1 may contribute to docetaxel resistance in PRAD patients (*p* = 0.018; **Figure 9, right**). A similar trend was observed in BRCA patients for cisplatin, although the difference did not reach statistical significance (*p* = 0.064; **Figure 9, left**). Taken together, these results showed that the involvement of SRPK1 in response to chemotherapeutic drugs is highly context-and cancer-type-dependent, indicating that the mechanisms of cisplatin and docetaxel resistance may be associated with altered SRPK1 expression, and highlighting that the drug-response prediction platform can identify clinically relevant genes affecting drug responses.

**Figure 9.**
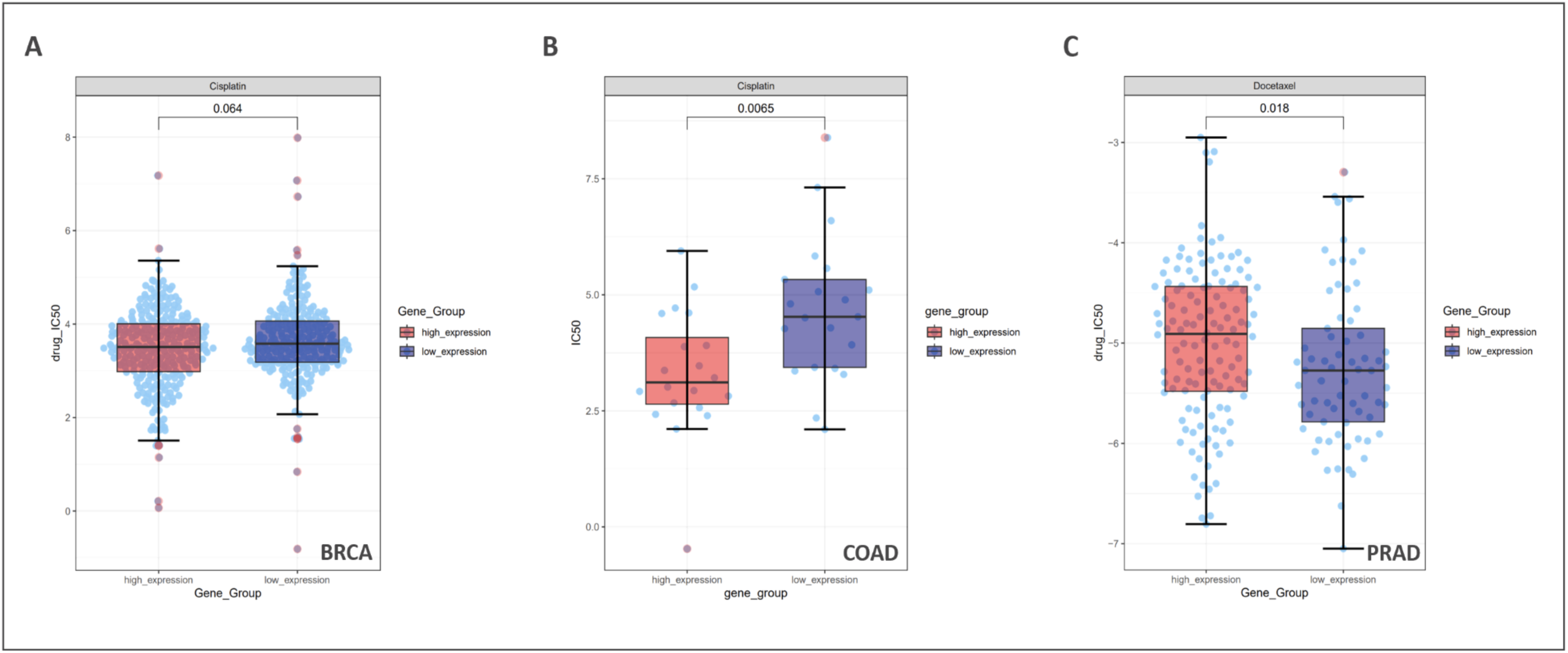
SRPK1 expression may predict sensitivity to chemotherapeutic agents in BRCA, COAD, and PRAD patient samples. Boxplots compares the predicted IC_50_ values of two chemotherapeutic agents, namely cisplatin and docetaxel, with SRPK1 mRNA expression, which were divided into SRPK1-high group and SRPK1-low group on the basis of drug sensitivity prediction from cancer-specific pharmacogenomics data of SRPK1 mRNA in tumour samples from the **(A)** TCGA-BRCA, **(B)** TCGA-COAD, and **(C)** TCGA-PRAD datasets analysed using the CPADS platform. The x-axis represents SRPK1 expression groups. The y-axis represents the predicted IC_50_ values of the chemotherapeutic agent within each of the groups. The colours on the legend boxes next to the plot represent high-SRPK1 expression (**light red**) and low-SRPK1 expression (**purple**) groups, respectively. Each circle (**light blue**) depicts the predicted IC_50_ value of an individual of the tumour sample. The middle line in the boxplot represents the median, with its lower and upper boundaries representing the first and third quartiles and whiskers extending to 1.5× the interquartile range of the quartiles. Two-sided Wilcoxon rank-sum test was conducted to determine the statistical significance of the difference between the two expression groups (*p* < 0.05 indicates statistical significance) significant.

## 4. Discussion

The complex and multifaceted challenge of chemoresistance necessitates the discovery of novel molecular targets and a deeper understanding of chemoresistance mechanisms in cancer cells. This study’s comprehensive pan-cancer analysis presented a robust, clinically-focused investigation into the role of SRPK1, a key regulator of alternative splicing, by integrating diverse multi-omics data.

Findings on the broad upregulation of SRPK1 mRNA across numerous tumour types, including significant elevation in BRCA and COAD at both mRNA and protein levels, strongly supported its role as a tumour driver. This was further substantiated by the prognostic analysis, which indicated the significant association of high SRPK1 expression, particularly in BRCA, with poor OS and DFS. In the context of cancer progression, genes associated with poor patient prognosis are often considered pivotal drivers and high-priority therapeutic targets due to their involvement in tumour-specific dependencies.

The functional association analysis, specifically the focus on the splicing factor SRSF1, provided a mechanistic link to the pathogenesis of chemoresistance. SRPK1 phosphorylates SR proteins like SRSF1, thereby regulating their activity and the downstream process of alternative splicing. Given that aberrant AS is an established mechanism of drug resistance—where changes in splicing factor abundance promote malignant progression and chemoresistance—the co-expression network of SRPK1 and SRSF1 suggests this pathway as a key player in tumour-specific malignancy. For instance, the AS of the pyruvate kinase gene (PKM) into its PKM2 isoform, regulated by the splicing factor PTBP1, has been associated with poor prognosis and drug resistance in pancreatic cancer, a parallel that highlights the significance of the SRPK1-SRSF1 axis in other cancers.

This study’s pharmacogenomics analysis was deemed crucial for translational relevance. The observed correlation between high SRPK1 expression and predicted resistance to agents like cisplatin and docetaxel in tumour cohorts provided strong clinical justification for targeting SRPK1. Findings reconciled the seemingly contradictory literature on SRPK1’s role in chemosensitivity by suggesting a tumour-specific and context-dependent modulation of the protein. For instance, while some studies found that SRPK1 downregulation restored cisplatin sensitivity, others suggested a link between reduced expression and enhanced resistance. The adopted multi-omics approach in this study—by defining the expression-to-prognosis-to-drug-response axis in a pan-cancer manner—introduced a more defined landscape where SRPK1-mediated effects are likely dependent on the specific tumour type and its core regulatory networks.

In conclusion, this study established SRPK1 as an overexpressed oncogenic factor associated with poor patient outcomes in key cancer types. The systematic evidence linking SRPK1 to a core splicing regulatory network (SRPK1-SRSF1) and predicted drug resistance pathways provided compelling evidence on SRPK1 as a promising therapeutic target for overcoming tumour-specific chemoresistance. Future experimental studies should validate the functional consequences of SRPK1 inhibition on the SRSF1-regulated spliceome in BRCA, COAD, and PRAD and assess the potential of SRPK1 inhibitors as adjuvants to the current chemotherapy regimens.

## Acknowledgments

The authors acknowledge the use of publicly available resources including TCGA, CPTAC, GEPIA2, and UALCAN.

## References

1. Aleksakhina, S.N., Kashyap, A. and Imyanitov, E.N. (2019) ‘Mechanisms of acquired tumor drug resistance’, Biochimica et Biophysica Acta (BBA) – Reviews on Cancer, 1872(2), p. 188310. Available at: 10.1016/J.BBCAN.2019.188310.

2. Anczukow, O. et al. (2024) ‘Steering research on mRNA splicing in cancer towards clinical translation’, Nature Reviews Cancer, 24(12), pp. 887–905. Available at: 10.1038/S41568-024-00750-2;SUBJMETA.

3. Avci, C.B. et al. (2025) ‘Precision oncology: Using cancer genomics for targeted therapy advancements’, Biochimica et Biophysica Acta (BBA) – Reviews on Cancer, 1880(1), p. 189250. Available at: 10.1016/J.BBCAN.2024.189250.

4. Bai, H. and Chen, B. (2020) ‘Abnormal PTBP1 Expression Sustains the Disease Progression of Multiple Myeloma’, Disease Markers, 2020(1), p. 4013658. Available at: 10.1155/2020/4013658.

5. Bian, Z. et al. (2024) ‘LINC01852 inhibits the tumorigenesis and chemoresistance in colorectal cancer by suppressing SRSF5-mediated alternative splicing of PKM’, Molecular Cancer, 23(1), pp. 1–17. Available at: 10.1186/S12943-024-01939-7/FIGURES/8.

6. Bray Bsc, F., et al. (2024) ‘Global cancer statistics 2022: GLOBOCAN estimates of incidence and mortality worldwide for 36 cancers in 185 countries’. Available at: 10.3322/caac.21834.

7. Brenner, D.R. et al. (2023) ‘Exploring the Future of Cancer Impact in Alberta: Projections and Trends 2020–2040’, Current Oncology 2023, Vol. 30, Pages 9981-9995, 30(11), pp. 9981–9995. Available at: 10.3390/CURRONCOL30110725.

8. Calabretta, S. et al. (2015) ‘Modulation of PKM alternative splicing by PTBP1 promotes gemcitabine resistance in pancreatic cancer cells’, Oncogene, 35(16), p. 2031. Available at: 10.1038/ONC.2015.270.

9. Campbell, P.J. et al. (2020) ‘Pan-cancer analysis of whole genomes’, Nature 2020 578:7793, 578(7793), pp. 82–93. Available at: 10.1038/s41586-020-1969-6.

10. Chen, C. et al. (2020) ‘LncRNA H19 downregulation confers erlotinib resistance through upregulation of PKM2 and phosphorylation of AKT in EGFR-mutant lung cancers’, Cancer Letters, 486, pp. 58–70. Available at: 10.1016/J.CANLET.2020.05.009.

11. Chen, F. et al. (2019) ‘Pan-cancer molecular subtypes revealed by mass-spectrometry-based proteomic characterization of more than 500 human cancers’, Nature Communications 2019 10:1, 10(1), pp. 1–15. Available at: 10.1038/s41467-019-13528-0.

12. Cheng, C. et al. (2018) ‘PTBP1 knockdown overcomes the resistance to vincristine and oxaliplatin in drug-resistant colon cancer cells through regulation of glycolysis’, Biomedicine & Pharmacotherapy, 108, pp. 194–200. Available at: 10.1016/J.BIOPHA.2018.09.031.

13. Cree, I.A. and Charlton, P. (2017) ‘Molecular chess? Hallmarks of anti-cancer drug resistance’, BMC Cancer, 17(1), pp. 1–8. Available at: 10.1186/S12885-016-2999-1/FIGURES/1.

14. Das, S. and Krainer, A.R. (2014) ‘Emerging functions of SRSF1, splicing factor and oncoprotein, in RNA metabolism and cancer’, Molecular Cancer Research, 12(9), pp. 1195–1204. Available at: 10.1158/1541-7786.MCR-14-0131/80622/AM/EMERGING-FUNCTIONS-OF-SRSF1-SPLICING-FACTOR-AND.

15. Deng, K. et al. (2021) ‘Abnormal alternative splicing promotes tumor resistance in targeted therapy and immunotherapy’, Translational Oncology, 14(6), p. 101077. Available at: 10.1016/J.TRANON.2021.101077.

16. Duzgun, D. and Oltean, S. (2025) ‘Aberrant Splicing as a Mechanism for Resistance to Cancer Therapies’, Cancers 2025, Vol. 17, Page 1381, 17(8), p. 1381. Available at: 10.3390/CANCERS17081381.

17. Fonseca-Montaño, M.A. et al. (2022) ‘Cancer Genomics’, Archives of Medical Research, 53(8), pp. 723–731. Available at: 10.1016/J.ARCMED.2022.11.011.

18. Goel, S. et al. (2016) ‘Overcoming Therapeutic Resistance in HER2-Positive Breast Cancers with CDK4/6 Inhibitors’, Cancer Cell, 29(3), pp. 255–269. Available at: 10.1016/j.ccell.2016.02.006.

19. Gong, L. et al. (2016) ‘Serine-arginine protein kinase 1 promotes a cancer stem cell-like phenotype through activation of Wnt/β-catenin signalling in NSCLC’, Journal of Pathology, 240(2), pp. 184–196. Available at: 10.1002/PATH.4767;JOURNAL:JOURNAL:15552039;WGROUP:STRING:PUBLICATION.

20. Huang, J.Q. et al. (2023) ‘Serine-arginine protein kinase 1 (SRPK1) promotes EGFR-TKI resistance by enhancing GSK3β Ser9 autophosphorylation independent of its kinase activity in non-small-cell lung cancer’, Oncogene 2023 42:15, 42(15), pp. 1233–1246. Available at: 10.1038/s41388-023-02645-2.

21. Hutter, C. and Zenklusen, J.C. (2018) ‘The Cancer Genome Atlas: Creating Lasting Value beyond Its Data’, Cell, 173(2), pp. 283–285. Available at: 10.1016/J.CELL.2018.03.042.

22. Jiang, P. et al. (2018) ‘Genome-Scale Signatures of Gene Interaction from Compound Screens Predict Clinical Efficacy of Targeted Cancer Therapies’, Cell Systems, 6(3), pp. 343–354.e5. Available at: 10.1016/j.cels.2018.01.009.

23. Jiang, P. et al. (2022) ‘Big data in basic and translational cancer research’, Nature Reviews Cancer 2022 22:11, 22(11), pp. 625–639. Available at: 10.1038/s41568-022-00502-0.

24. Kaelin, W.G. (2017) ‘Common pitfalls in preclinical cancer target validation’, Nature Reviews Cancer, 17(7), pp. 441–450. Available at: 10.1038/NRC.2017.32;SUBJMETA=1059,153,154,631,67,70;KWRD=CANCER+MODELS,CANCER+THERAPY,DRUG+DEVELOPMENT,DRUG+DISCOVERY.

25. Krishnakumar, S. et al. (2008) ‘SRPK1: A cisplatin sensitive protein expressed in retinoblastoma’, Pediatric Blood and Cancer, 50(2), pp. 402–406. Available at: 10.1002/PBC.21088;JOURNAL:JOURNAL:1096911X;REQUESTEDJOURNAL:JOURNAL:15455017;WGROUP:STRING:PUBLICATION.

26. Li, K. et al. (2024) ‘CPADS: a web tool for comprehensive pancancer analysis of drug sensitivity’, Briefings in Bioinformatics, 25(3). Available at: 10.1093/BIB/BBAE237.

27. Li, W. et al. (2024) ‘SNRNP70 regulates the splicing of CD55 to promote osteosarcoma progression’, JCI Insight, 9(24). Available at: 10.1172/JCI.INSIGHT.185269.

28. Liu, B. et al. (2024) ‘Exploring treatment options in cancer: tumor treatment strategies’, Signal Transduction and Targeted Therapy 2024 9:1, 9(1), pp. 1–44. Available at: 10.1038/s41392-024-01856-7.

29. Liu, C.J. et al. (2018) ‘GSCALite: a web server for gene set cancer analysis’, Bioinformatics, 34(21), pp. 3771–3772. Available at: 10.1093/BIOINFORMATICS/BTY411.

30. Liu, H. et al. (2021) ‘SRPK1/2 and PP1α exert opposite functions by modulating SRSF1-guided MKNK2 alternative splicing in colon adenocarcinoma’, Journal of Experimental and Clinical Cancer Research, 40(1), pp. 1–16. Available at: 10.1186/S13046-021-01877-Y/FIGURES/7.

31. Liu, Y. et al. (2017) ‘Impact of Alternative Splicing on the Human Proteome’, Cell Reports, 20(5), p. 1229. Available at: 10.1016/J.CELREP.2017.07.025.

32. López-Cánovas, J.L. et al. (2023) ‘PRPF8 increases the aggressiveness of hepatocellular carcinoma by regulating FAK/AKT pathway via fibronectin 1 splicing’, Experimental & Molecular Medicine 2023 55:1, 55(1), pp. 132–142. Available at: 10.1038/s12276-022-00917-7.

33. Lv, Y. et al. (2019) ‘Erlotinib overcomes paclitaxel-resistant cancer stem cells by blocking the EGFR-CREB/GRβ-IL-6 axis in MUC1-positive cervical cancer’, Oncogenesis, 8(12), pp. 1–12. Available at: 10.1038/S41389-019-0179-2;TECHMETA.

34. Madukwe, J.C. (2023) ‘Overcoming drug resistance in cancer’, Cell, 186(8), pp. 1515–1516. Available at: 10.1016/j.cell.2023.03.019.

35. Malvi, P. et al. (2020) ‘LIMK2 promotes the metastatic progression of triple-negative breast cancer by activating SRPK1’, Oncogenesis, 9(8), pp. 1–16. Available at: 10.1038/S41389020002631;TECHMETA=106,58,82,96;SUBJMETA=1347,208,631,67,68;KWRD=BREAST+CANCER,CANCER+GENETICS.

36. Marusyk, A., Janiszewska, M. and Polyak, K. (2020) ‘Intratumor Heterogeneity: The Rosetta Stone of Therapy Resistance’, Cancer Cell, 37(4), pp. 471–484. Available at: 10.1016/J.CCELL.2020.03.007.

37. Mavrou, A. et al. (2014) ‘Serine–arginine protein kinase 1 (SRPK1) inhibition as a potential novel targeted therapeutic strategy in prostate cancer’, Oncogene 2015 34:33, 34(33), pp. 4311–4319. Available at: 10.1038/onc.2014.360.

38. Mehterov, N. et al. (2021) ‘Alternative RNA Splicing—The Trojan Horse of Cancer Cells in Chemotherapy’, Genes 2021, Vol. 12, Page 1085, 12(7), p. 1085. Available at: 10.3390/GENES12071085.

39. Nagai, H. and Kim, Y.H. (2017) ‘Cancer prevention from the perspective of global cancer burden patterns’, Journal of Thoracic Disease, 9(3), p. 448. Available at: 10.21037/JTD.2017.02.75.

40. Nikas, I.P. et al. (2019) ‘Serine-Arginine Protein Kinase 1 (SRPK1) as a Prognostic Factor and Potential Therapeutic Target in Cancer: Current Evidence and Future Perspectives’, Cells 2020, Vol. 9, Page 19, 9(1), p. 19. Available at: 10.3390/CELLS9010019.

41. Odunsi, K. et al. (2012) ‘Elevated Expression of the Serine-Arginine Protein Kinase 1 Gene in Ovarian Cancer and Its Role in Cisplatin Cytotoxicity In Vitro’, PLOS ONE, 7(12), p. e51030. Available at: 10.1371/JOURNAL.PONE.0051030.

42. Pal, A.S. et al. (2021) ‘Identification of microRNAs that promote erlotinib resistance in non-small cell lung cancer’, Biochemical Pharmacology, 189, p. 114154. Available at: 10.1016/J.BCP.2020.114154.

43. Pan, R. and Zhou, H. (2020) ‘Exosomal Transfer of lncRNA H19 Promotes Erlotinib Resistance in Non-Small Cell Lung Cancer via miR-615-3p/ATG7 Axis’. Available at: 10.2147/CMAR.S241095.

44. Pan, Z. et al. (2023) ‘Role of NAT10-mediated ac4C-modified HSP90AA1 RNA acetylation in ER stress-mediated metastasis and lenvatinib resistance in hepatocellular carcinoma’, Cell Death Discovery, 9(1), pp. 1–14. Available at: 10.1038/S41420-023-01355-8;SUBJMETA=1275,2327,458,631,67,80;KWRD=ACETYLATION,CANCER+METABOLISM.

45. Patel, A.S. and Yanai, I. (2024) ‘A developmental constraint model of cancer cell states and tumor heterogeneity’, Cell, 187(12), pp. 2907–2918. Available at: 10.1016/J.CELL.2024.04.032.

46. Patel, M., Sachidanandan, M. and Adnan, M. (2018) ‘Serine arginine protein kinase 1 (SRPK1): a moonlighting protein with theranostic ability in cancer prevention’, Molecular Biology Reports 2018 46:1, 46(1), pp. 1487–1497. Available at: 10.1007/S11033-018-4545-5.

47. Pellarin, I. et al. (2020) ‘Splicing factor proline– and glutamine-rich (SFPQ) protein regulates platinum response in ovarian cancer-modulating SRSF2 activity’, Oncogene 2020 39:22, 39(22), pp. 4390–4403. Available at: 10.1038/s41388-020-1292-6.

48. Qi, Y. et al. (2023) ‘Cuproptosis-related gene SLC31A1: prognosis values and potential biological functions in cancer’, Scientific Reports, 13(1), pp. 1–14. Available at: 10.1038/S41598-023-44681-8;SUBJMETA=114,631,67;KWRD=CANCER,COMPUTATIONAL+BIOLOGY+AND+BIOINFORMATICS.

49. Qian, J. et al. (2020) ‘A pan-cancer blueprint of the heterogeneous tumor microenvironment revealed by single-cell profiling’, Cell Research 2020 30:9, 30(9), pp. 745–762. Available at: 10.1038/s41422-020-0355-0.

50. Ramos, A., Sadeghi, S. and Tabatabaeian, H. (2021) ‘Battling Chemoresistance in Cancer: Root Causes and Strategies to Uproot Them’, International Journal of Molecular Sciences, 22(17), p. 9451. Available at: 10.3390/IJMS22179451.

51. Ren, Y. et al. (2022) ‘GBP2 facilitates the progression of glioma via regulation of KIF22/EGFR signaling’, Cell Death Discovery, 8(1), pp. 1–9. Available at: 10.1038/S41420-022-01018-0;SUBJMETA=395,631,67,68,692,699;KWRD=CANCER+GENETICS,ONCOGENES.

52. Roy, R. et al. (2019) ‘Expression levels of therapeutic targets as indicators of sensitivity to targeted therapeutics’, Molecular Cancer Therapeutics, 18(12), pp. 2480–2489. Available at: 10.1158/1535-7163.MCT-19-0273/87988/AM/EXPRESSION-LEVELS-OF-THERAPEUTIC-TARGETS-AS.

53. Ruprecht, B. et al. (2017) ‘Lapatinib resistance in breast cancer cells is accompanied by phosphorylation-mediated reprogramming of glycolysis’, Cancer Research, 77(8), pp. 1842–1853. Available at: 10.1158/0008-5472.CAN-16-2976/652733/AM/LAPATINIB-RESISTANCE-IN-BREAST-CANCER-CELLS-IS.

54. Samantaray, A. et al. (2024) ‘Novel insight into cancer treatment: Recent advances and new challenges’, Journal of Drug Delivery Science and Technology, 93, p. 105384. Available at: 10.1016/J.JDDST.2024.105384.

55. Schenk, P.W. et al. (2001) ‘SKY1 Is Involved in Cisplatin-induced Cell Kill in Saccharomyces cerevisiae, and Inactivation of Its Human Homologue, SRPK1, Induces Cisplatin Resistance in a Human Ovarian Carcinoma Cell Line 1’, CANCER RESEARCH, 61, pp. 6982–6986. Available at: http://aacrjournals.org/cancerres/article-pdf/61/19/6982/2487357/ch1901006982.pdf (Accessed: 18 June 2025).

56. Shea, A. et al. (2025) ‘Modeling Drug Responses and Evolutionary Dynamics Using Patient-Derived Xenografts Reveals Precision Medicine Strategies for Triple-Negative Breast Cancer’, Cancer Research, 85(3), pp. 567–584. Available at: 10.1158/0008-5472.CAN-24-1703/749968/AM/MODELING-DRUG-RESPONSES-AND-EVOLUTIONARY-DYNAMICS.

57. Sheng, J. et al. (2018) ‘SRSF1 modulates PTPMT1 alternative splicing to regulate lung cancer cell radioresistance’, eBioMedicine, 38, pp. 113–126. Available at: 10.1016/J.EBIOM.2018.11.007.

58. Sinnakannu, J.R. et al. (2020) ‘SRSF1 mediates cytokine-induced impaired imatinib sensitivity in chronic myeloid leukemia’, Leukemia 2020 34:7, 34(7), pp. 1787–1798. Available at: 10.1038/s41375-020-0732-1.

59. Steggall, J. et al. (2025) ‘Integrative proteo-genomic profiling uncovers key biomarkers of lapatinib resistance in HER2-positive breast cancer’, British Journal of Cancer, pp. 1–12. Available at: 10.1038/S41416-025-03174-3;SUBJMETA.

60. Sveen, A. et al. (2016) ‘Aberrant RNA splicing in cancer; Expression changes and driver mutations of splicing factor genes’, Oncogene, 35(19), pp. 2413–2427. Available at: 10.1038/ONC.2015.318;SUBJMETA=208,631,69;KWRD=CANCER+GENOMICS.

61. Szklarczyk, D. et al. (2015) ‘STRING v10: protein–protein interaction networks, integrated over the tree of life’, Nucleic Acids Research, 43(D1), pp. D447–D452. Available at: 10.1093/NAR/GKU1003.

62. Tang, Z. et al. (2017) ‘GEPIA: a web server for cancer and normal gene expression profiling and interactive analyses’, Nucleic Acids Research, 45(W1), pp. W98–W102. Available at: 10.1093/NAR/GKX247.

63. Van Roosmalen, W. et al. (2015) ‘Tumor cell migration screen identifies SRPK1 as breast cancer metastasis determinant’, The Journal of Clinical Investigation, 125(4), pp. 1648–1664. Available at: 10.1172/JCI74440.

64. Vokes, N.I. et al. (2022) ‘Concurrent TP53 Mutations Facilitate Resistance Evolution in EGFR-Mutant Lung Adenocarcinoma’, Journal of Thoracic Oncology, 17(6), pp. 779–792. Available at: 10.1016/J.JTHO.2022.02.011.

65. Wagner, K.D. et al. (2019) ‘Altered VEGF Splicing Isoform Balance in Tumor Endothelium Involves Activation of Splicing Factors Srpk1 and Srsf1 by the Wilms’ Tumor Suppressor Wt1’, Cells 2019, Vol. 8, Page 41, 8(1), p. 41. Available at: 10.3390/CELLS8010041.

66. Wang, C. et al. (2020) ‘SRPK1 acetylation modulates alternative splicing to regulate cisplatin resistance in breast cancer cells’, Communications Biology 2020 3:1, 3(1), pp. 1–13. Available at: 10.1038/s42003-020-0983-4.

67. Wang, Z. et al. (2012) ‘Exon-centric regulation of pyruvate kinase M alternative splicing via mutually exclusive exons’, Journal of Molecular Cell Biology, 4(2), pp. 79–87. Available at: 10.1093/JMCB/MJR030.

68. Wang, Z.W. et al. (2022) ‘SRSF3-mediated regulation of N6-methyladenosine modification-related lncRNA ANRIL splicing promotes resistance of pancreatic cancer to gemcitabine’, Cell Reports, 39(6), p. 110813. Available at: 10.1016/J.CELREP.2022.110813/ATTACHMENT/3A87CD8A-12CE-429F-B0E1-FD9B428A8CC0/MMC2.XLSX.

69. Warde-Farley, D. et al. (2010) ‘The GeneMANIA prediction server: biological network integration for gene prioritization and predicting gene function’, Nucleic Acids Research, 38(Web Server issue), p. W214. Available at: 10.1093/NAR/GKQ537.

70. Wu, G. et al. (2021) ‘ZNF711 down-regulation promotes CISPLATIN resistance in epithelial ovarian cancer via interacting with JHDM2A and suppressing SLC31A1 expression’, eBioMedicine, 71, p. 103558. Available at: 10.1016/J.EBIOM.2021.103558.

71. Xie, R. et al. (2023) ‘NAT10 Drives Cisplatin Chemoresistance by Enhancing ac4C-Associated DNA Repair in Bladder Cancer’, Cancer Research, 83(10), pp. 1666–1683. Available at: 10.1158/0008-5472.CAN-22-2233/718819/AM/NAT10-DRIVES-CISPLATIN-CHEMORESISTANCE-BY.

72. Yamasaki, F. et al. (2007) ‘Sensitivity of breast cancer cells to erlotinib depends on cyclin-dependent kinase 2 activity’, Molecular Cancer Therapeutics, 6(8), pp. 2168–2177. Available at: 10.1158/1535-7163.MCT-06-0514/357316/P/SENSITIVITY-OF-BREAST-CANCER-CELLS-TO-ERLOTINIB.

73. Yang, Y. et al. (2022) ‘Study on the prognosis, immune and drug resistance of m6A-related genes in lung cancer’, BMC Bioinformatics, 23(1), pp. 1–23. Available at: 10.1186/S12859-022-04984-5/FIGURES/1.

74. Yi, N. et al. (2018) ‘SRPK1 is a poor prognostic indicator and a novel potential therapeutic target for human colorectal cancer’, OncoTargets and therapy, 11, p. 5359. Available at: 10.2147/OTT.S172541.

75. Yi, X. et al. (2023) ‘SIRT7 orchestrates melanoma progression by simultaneously promoting cell survival and immune evasion via UPR activation’, Signal Transduction and Targeted Therapy, 8(1), pp. 1–17. Available at: 10.1038/S41392-023-01314W;SUBJMETA=1813,327,631,67;KWRD=CANCER+MICROENVIRONMENT,SKIN+CANCER.

76. Zahra, K. et al. (2020) ‘Pyruvate Kinase M2 and Cancer: The Role of PKM2 in Promoting Tumorigenesis’, Frontiers in Oncology, 10, p. 505842. Available at: 10.3389/FONC.2020.00159/BIBTEX.

77. Zhang, F. et al. (2021) ‘LncRNA CRNDE attenuates chemoresistance in gastric cancer via SRSF6-regulated alternative splicing of PICALM’, Molecular Cancer, 20(1), pp. 1–6. Available at: 10.1186/S12943-020-01299-Y/FIGURES/3.

78. Zhao, C. et al. (2021) ‘Splicing Factor DDX23, Transcriptionally Activated by E2F1, Promotes Ovarian Cancer Progression by Regulating FOXM1’, Frontiers in Oncology, 11, p. 749144. Available at: 10.3389/FONC.2021.749144/BIBTEX.

