## Supplementary files for "Clinical Implications of SRPK1 Expression in Human Tumours: A Comprehensive Pan-Cancer Analysis Based on Multi-Omics Databases"

### Appendix 1: Supplemental Information

**Supplementary Table S1: SRPK1 exhibits frequent differential expression across TCGA tumours, with alterations classified as up (upregulation), down (downregulation), or n.s. (not significant) based on gene expression levels.**

| Cancer type | Disease full name | SRPK1 mRNA expression in cancer |
|---|---|---|
| BLCA | Bladder urothelial carcinoma | Up |
| BRCA | Breast invasive carcinoma | Up |
| CESC | Cervical squamous cell carcinoma and endocervical carcinoma | Up |
| CHOL | Cholangiocarcinoma | Up |
| COAD | Colon adenocarcinoma | Up |
| ESCA | Esophageal carcinoma | Up |
| HNSC | Head and neck squamous cell carcinoma | Up |
| LIHC | Liver hepatocellular carcinoma | Up |
| LUAD | Lung adenocarcinoma | Up |
| LUSC | Lung squamous cell carcinoma | Up |
| READ | Rectum adenocarcinoma | Up |
| STAD | Stomach adenocarcinoma | Up |
| UCEC | Uterine corpus endometrial carcinoma | Up |
| KICH | Kidney chromophobe | Down |
| KIRC | Kidney renal clear cell carcinoma | Down |
| GBM | Glioblastoma multiforme | n.s. |
| KIRP | Kidney renal papillary cell carcinoma | n.s. |
| PAAD | Pancreatic adenocarcinoma | n.s. |
| PRAD | Prostate adenocarcinoma | n.s. |
| THCA | Thyroid carcinoma | n.s. |

BRCA, COAD, and PRAD are highlighted in red.

**Supplementary Table S2: The description of SRPK1-related top 28 genes identified by analysing GeneMANIA and STRING databases (Related to Figure 6).**

| Gene | Full Name | Source (NCBI) | Database |
|---|---|---|---|
| DDX23 | DEAD-box helicase 23 | https://www.ncbi.nlm.nih.gov/gene/9416 | STRING |
| GBP2 | Guanylate binding protein 2 | https://www.ncbi.nlm.nih.gov/gene/2634 | GeneMANIA |
| GTF3C4 | General transcription factor IIIC subunit 4 | https://www.ncbi.nlm.nih.gov/gene/9329 | GeneMANIA |
| HECW2 | HECT, C2, and WW domain containing E3 ubiquitin protein ligase 2 | https://www.ncbi.nlm.nih.gov/gene/57520 | GeneMANIA |
| LBR | Lamin B receptor | https://www.ncbi.nlm.nih.gov/gene/3930 | GeneMANIA |
| NAT10 | N-acetyltransferase 10 | https://www.ncbi.nlm.nih.gov/gene/55226 | GeneMANIA |
| PPM1G | Protein phosphatase, Mg2+/Mn2+ dependent 1G | https://www.ncbi.nlm.nih.gov/gene/5496 | GeneMANIA |
| PRM1 | Protamine 1 | https://www.ncbi.nlm.nih.gov/gene/5619 | GeneMANIA |
| PRPF38A | Pre-mRNA processing factor 38A | https://www.ncbi.nlm.nih.gov/gene/84950 | GeneMANIA |
| PRPF8 | Pre-mRNA processing factor 8 | https://www.ncbi.nlm.nih.gov/gene/10594 | GeneMANIA |
| RBM4 | RNA binding motif protein 4 | https://www.ncbi.nlm.nih.gov/gene/5936 | GeneMANIA |
| SAFB | Scaffold attachment factor B | https://www.ncbi.nlm.nih.gov/gene/6294 | GeneMANIA |
| SAFB2 | Scaffold attachment factor B2 | https://www.ncbi. nlm.nih.gov/gene/9667 | GeneMANIA |
| SERPINB10 | Serpin family B member 10 | https://www.ncbi.nlm.nih.gov/gene/5273 | GeneMANIA |

| | | | |
|---|---|---|---|
| SF3A3 | Splicing factor 3a subunit 3 | https://www.ncbi.nlm.nih.gov/gene/10946 | GeneMANIA |
| SIRT7 | Sirtuin 7 | https://www.ncbi.nlm.nih.gov/gene/51547 | GeneMANIA |
| SNRNP70 | Small nuclear ribonucleoprotein U1 subunit 70 | https://www.ncbi.nlm.nih.gov/gene/6625 | STRING |
| SRSF1 | Serine and arginine rich splicing factor 1 | https://www.ncbi.nlm.nih.gov/gene/6426 | GeneMANIA STRING |
| SRSF2 | Serine and arginine rich splicing factor 2 | https://www.ncbi.nlm.nih.gov/gene/6427 | STRING |
| SRSF3 | Serine and arginine rich splicing factor 3 | https://www.ncbi.nlm.nih.gov/gene/6428 | STRING |
| SRSF4 | Serine and arginine rich splicing factor 4 | https://www.ncbi.nlm.nih.gov/gene/6429 | STRING |
| SRSF5 | Serine and arginine rich splicing factor 5 | https://www.ncbi.nlm.nih.gov/gene/6430 | STRING |
| SRSF6 | Serine and arginine rich splicing factor 6 | https://www.ncbi.nlm.nih.gov/gene/6431 | GeneMANIA STRING |
| SRSF7 | Serine and arginine rich splicing factor 7 | https://www.ncbi.nlm.nih.gov/gene/6432 | STRING |
| SRSF9 | Serine and arginine rich splicing factor 9 | https://www.ncbi.nlm.nih.gov/gene/8683 | STRING |
| TARDBP | TAR DNA binding protein | https://www.ncbi.nlm.nih.gov/gene/23435 | GeneMANIA |
| ZRSR2 | Zinc finger CCH-type binding motif and serine/arginine rich 2 | https://www.ncbi.nlm.nih.gov/gene/8233 | GeneMANIA |

NCBI denotes National Centre for Biotechnology Information; GeneMANIA denotes Gene Multiple Association Network Integration Algorithm; STRING denotes Search Tool for the Retrieval of Integrating Proteins. SRSF1 is highlighted in red.

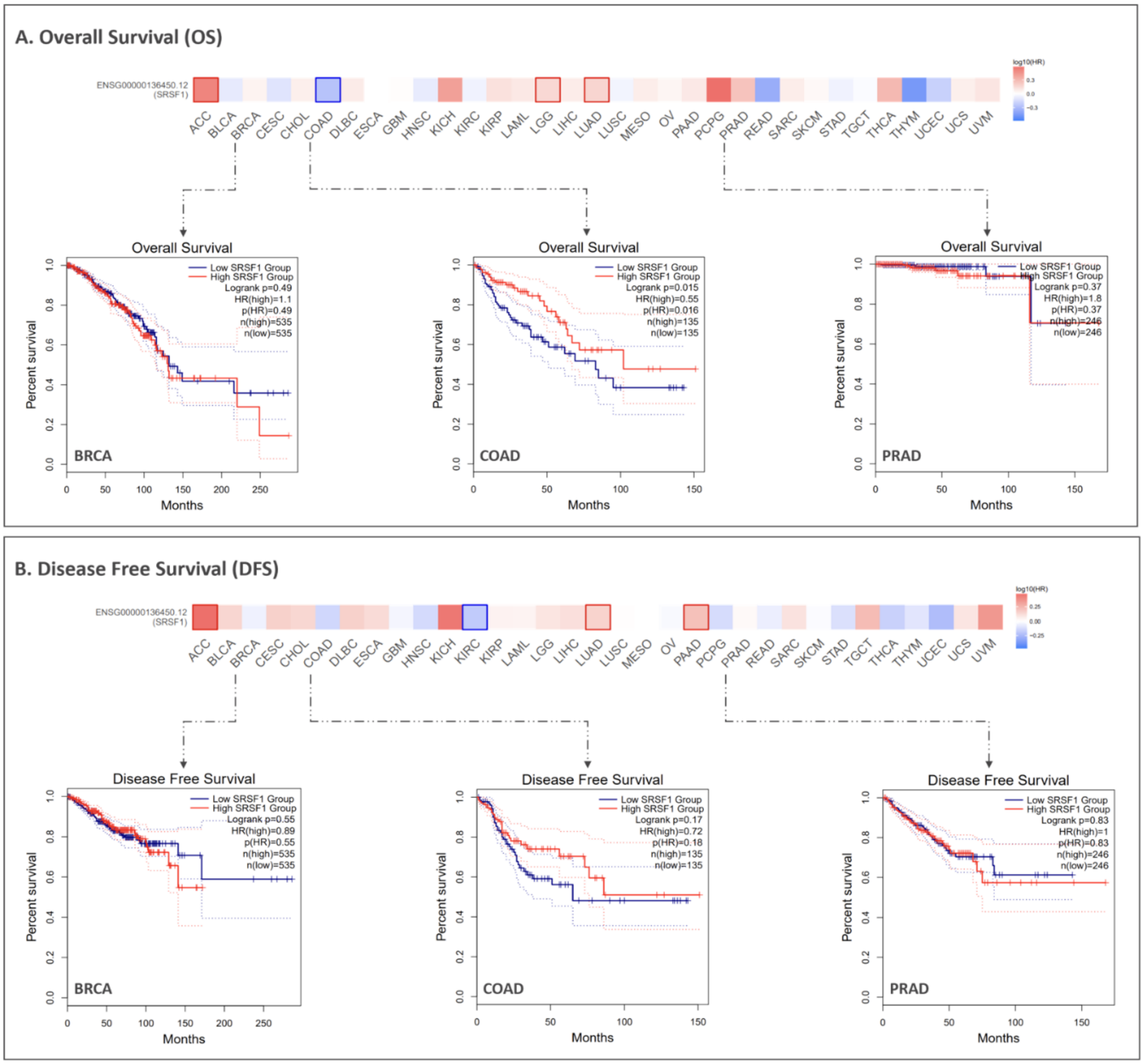


**Supplementary Figure S1: Prognostic implications of the selected target gene SRSF1 across BRCA, COAD, and PRAD clinical cohorts (Related to Figure 6). (A)** OS and **(B)** DFS analysed in BRCA, COAD, and PRAD patient cohorts using the GEPIA2 database. Patients in the indicated cancer cohorts were stratified based on the median cutoff point of SRSF1 mRNA expression to generate the Kaplan-Meier curves of OS and DFS (plotted in months). The **red** and **blue** lines represent the survival curves of patients with high and low levels of SRSF1 mRNA. Indicated by dashed lines, HR values (with 95% confidence intervals) represent the relative risk of the low-expression group compared with the high-expression group. HR > 1 implies the gene as a potential risk factor, whereas HR < 1 suggests a protective factor. Based on a log-rank (Mantel-Cox) test, a log-rank $p < 0.05$ indicates statistical significance. n indicates the patient count in each expression group.

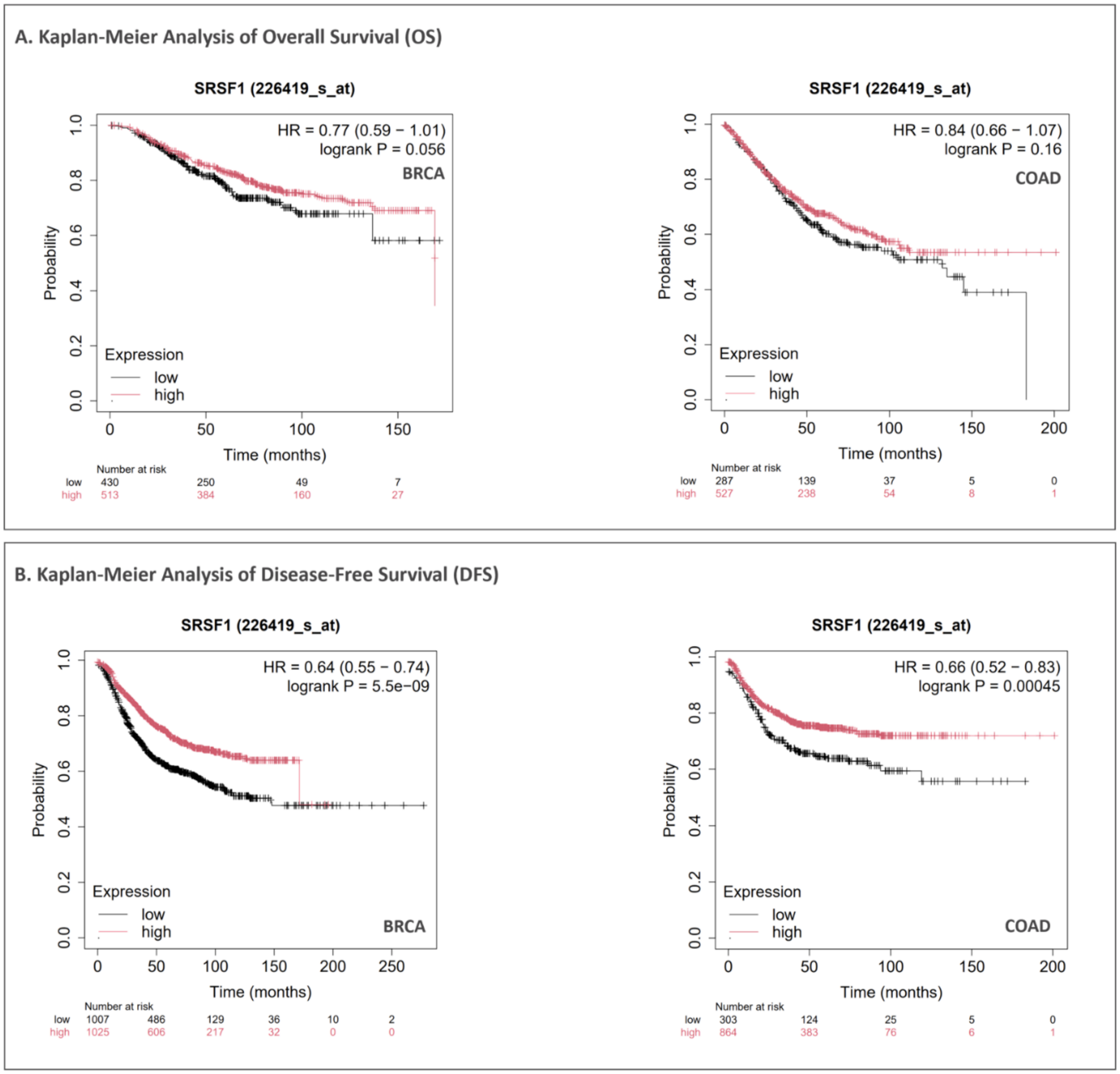


**Supplementary Figure S2: Prognostic relevance of the selected target gene SRSF1 in BRCA and COAD patients (Related to Figure 7).** Kaplan-Meier plots revealed differences in **(A)** OS and **(B)** DFS of BRCA and COAD patients based on the 226419_s_at probe, obtained from the Kaplan-Meier plotter database. The OS and DFS of BRCA and COAD patients stratified by low (**black** line) and high (**red** line) SRSF1 expression based on the optimal cutoff point provided by Kaplan-Meier plotter database. The OS and DFS of high- and low-expression groups were compared based on HR (with 95% confidence intervals). Based on a log-rank (Mantel-Cox) test, a log-rank $p < 0.05$ indicates statistical significance.

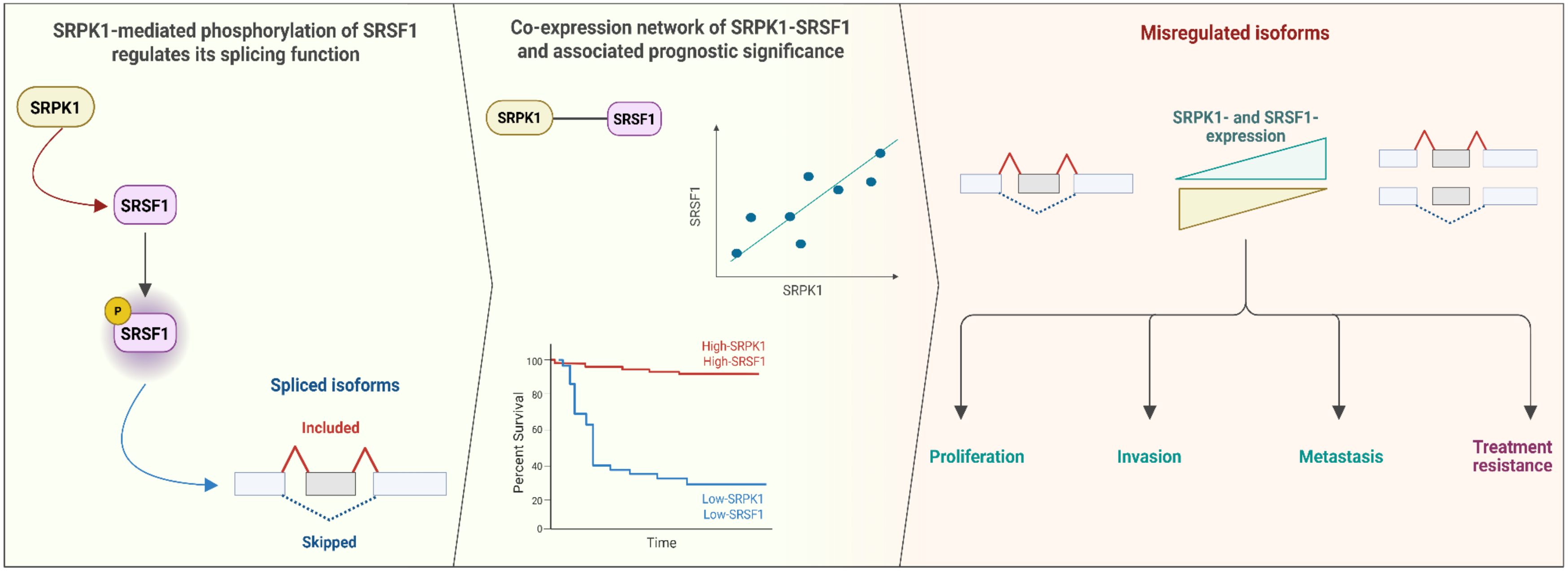


**Supplementary Figure S3: A proposed model of SRPK1-dependent SRSF1 expression that may drive alternative splicing isoform changes leading to oncogenic effects in a cancer-specific manner. (Left)** SRPK1 phosphorylates the splicing factor SRSF1 to regulate its functional activity in pre-mRNA splicing, thereby generating mRNA variants through the inclusion or skipping of exons or introns during the splicing process. Light blue boxes indicate exons, while a light grey box indicates an intron. Dotted **blue** lines connecting exons represent the skipped intron, and **red** lines represent the intron inclusion. P denotes phosphorylation. **(Middle)** Co-expression analysis between SRPK1 and SRSF1 at the mRNA level may identify cancer-specific biological roles with potentially crucial contributions to cancer progression across multiple cancer types. Independent survival analyses of SRPK1 and SRSF1 are likely to reveal crucial prognostic associations, whereby their expression changes correlate with favourable (or poor) outcomes in a cancer-specific manner. **(Right)** Co-upregulation (or co-downregulation) of SRPK1, together with SRSF1, may lead to changes in splicing isoforms, correlated with an increased probability of exon inclusion or skipping. Hence, in cancer cells, such distinct splicing isoforms may be involved in each of the steps of proliferation, invasion, metastasis, and treatment resistance, either directly or indirectly, through downstream targets. The **cyan** triangle represents increasing expression levels, and the **yellow** inverted triangle represents decreasing expression levels. Created with BioRender.com.

**Supplementary Table S3: List of experimental profiled anti-cancer agents showing the top nine positive and the top 21 negative correlations between drug response and SRPK1-high expression levels across diverse cancer cell lines in the GDSC dataset (Related to Figure 8A).**

| Anti-cancer agents | Correlation coefficient (*r*) | FDR | Pathway |
|---|---|---|---|
| 17-AAG | 0.170 | 2.44E-06 | Protein stability and degradation |
| CCT007093 | 0.175 | 8.19E-06 | Cell cycle |
| Erlotinib | 0.230 | 0.000246 | EGFR signalling |
| Lapatinib | 0.214 | 0.000207 | EGFR signalling |
| PD-0325901 | 0.172 | 3.79E-06 | ERK/MAPK signalling |
| PLX4720 | 0.161 | 9.23E-06 | ERK/MAPK signalling |
| RDEA119 | 0.185 | 7.66E-08 | ERK/MAPK signalling |
| Selumetinib | 0.185 | 7.78E-08 | ERK/MAPK signalling |
| TGX221 | 0.243 | 2.05E-05 | PIK3/mTOR signalling |
| 5-Fluorouracil | − 0.154 | 1.18E-05 | Other |
| AICAR | − 0.168 | 7.06E-06 | Metabolism |
| AR-42 | − 0.189 | 4.69E-08 | Chromatin histone acetylation |
| AT-7519 | − 0.209 | 9.41E-10 | Cell cycle |
| CP466722 | − 0.151 | 1.32E-05 | Genome integrity |
| GSK1070916 | − 0.172 | 9.89E-07 | Mitosis |
| I-BET-762 | − 0.188 | 3.35E-08 | Chromatin other |
| Ispinesib Mesylate | − 0.199 | 8.58E-09 | Mitosis |
| Methotrexate | − 0.193 | 5.8E-08 | DNA replication |
| Navitoclax | − 0.175 | 1.07E-06 | Apoptosis regulation |
| NPK76-II-72-1 | − 0.266 | 1.87E-15 | Cell cycle |
| PHA-793887 | − 0.227 | 1.9E-11 | Cell cycle |
| PIK-93 | − 0.156 | 5.78E-06 | PIK3/mTOR signalling |
| SNX-2112 | − 0.154 | 1.08E-05 | Protein stability and degradation |
| TAK-715 | − 0.165 | 2.1E-06 | JNK and p38 signalling |
| THZ-2-102-1 | − 0.185 | 9.58E-08 | Cell cycle |
| TPCA-1 | − 0.174 | 3.93E-07 | Kinases other |
| Tubastatin A | − 0.153 | 1.05E-05 | Chromatin histone acetylation |

| | | | |
|---|---|---|---|
| Vorinostat | − 0.201 | 1.47E-08 | Chromatin histone acetylation |
| WZ3105 | − 0.208 | 1.06E-09 | p53 signalling |
| XMD13-2 | − 0.174 | 4.82E-07 | Cytoskeleton |

R values represent Pearson’s correlation coefficients; the statistical significance of the correlation is determined based on the corresponding false discovery rate (FDR ≤ 0.05); r > 0 indicates positive correlation; r < 0 indicates negative correlation; erlotinib, lapatinib, and TGX221 are highlighted in red; AT-7519, NPK76-II-72-1, and PHA-793887 are highlighted in blue.

**Supplementary Table S4: List of experimental profiled anti-cancer agents showing the top 30 negative correlations between drug response and SRPK1-high expression levels across diverse cancer cell lines in the CTRP dataset (Related to Figure 8B).**

| Anti-cancer agents | Correlation coefficient (*r*) | FDR | Pathway |
|---|---|---|---|
| BI-2536 | − 0.32 | 4.15E-19 | Cell cycle |
| BRD-K66453893 | − 0.32 | 5.05E-18 | ERK/MAPK signalling other |
| BRD-K70511574 | − 0.34 | 3.59E-21 | Cell cycle |
| CD-437 | − 0.32 | 3.76E-18 | Apoptosis regulation other |
| CHM-1 | − 0.31 | 7.96E-18 | Mitosis |
| Ciclopirox | − 0.31 | 5.27E-18 | Chromatin histone acetylation other |
| COL-3 | − 0.38 | 2.16E-18 | Other |
| Cytarabine hydrochloride | − 0.31 | 2.41E-17 | p53 signalling other |
| Decitabine | − 0.31 | 1.66E-17 | Other |
| Fluorouracil | − 0.31 | 1.82E-17 | Metabolism other |
| FQI-1 | − 0.34 | 9.14E-11 | Other |
| FQI-2 | − 0.31 | 1.46E-17 | Other |
| GSK461364 | − 0.37 | 5.27E-25 | Cell cycle |
| Isoevodiamine | − 0.32 | 1.72E-18 | Apoptosis regulation other |
| KX2-391 | − 0.31 | 2.14E-17 | Metabolism other |
| Leptomycin B | − 0.35 | 3.36E-23 | p53 signalling |
| MK-1775 | − 0.34 | 9.85E-20 | Cell cycle |
| ML311 | − 0.31 | 6.96E-18 | Apoptosis regulation |
| Nakiterpiosin | − 0.33 | 1.74E-19 | Mitosis |
| Narciclasine | − 0.33 | 1.38E-19 | Other |
| NVP-231 | − 0.35 | 4.62E-22 | Other |
| Parbendazole | − 0.31 | 1.54E-18 | Mitosis |
| PHA-793887 | − 0.33 | 1.01E-19 | Cell cycle |
| Rigosertib | − 0.31 | 7.80E-17 | PIK3/mTOR signalling |
| SB-743921 | − 0.32 | 2.45E-19 | Mitosis |
| SR-II-138A | − 0.34 | 2.05E-21 | Other |
| Teniposide | − 0.34 | 3.94E-11 | DNA replication |
| Triazolothiadiazine | − 0.33 | 5.21E-20 | Other |

| | | | |
|---|---|---|---|
| Vincristine | − 0.32 | 4.26E-20 | Cytoskeleton |
| Vorinostat | − 0.31 | 6.17E-17 | Chromatin histone acetylation |

R values represent Pearson's correlation coefficients; the statistical significance of the correlation is determined based on the corresponding false discovery rate (FDR ≤ 0.05); r < 0 indicates negative correlation.
